# DNASE1L3 surveils mitochondrial DNA on the surface of distinct mammalian cells

**DOI:** 10.1101/2025.10.05.680464

**Authors:** Jennifer Porat, Valentina Poli, Karim Almahayni, Benson M. George, Francisca Nathália de Luna Vitorino, Li Yi, Carolina Bras Costa, Yitong Zhou, Andrew Crompton, Hongxiu Wen, Leon Zheng, Peter K. Gregersen, Suneet Agarwal, Benjamin A. Garcia, Yixuan Xie, Leonhard Möckl, Ivan Zanoni, Kim R. Simpfendorfer, Ryan A. Flynn

## Abstract

The extracellular space is a critical environment for discriminating self versus non-self nucleic acids and initiating the appropriate immune responses through signaling cascades to relay information about extracellular nucleic acids. Here, we provide evidence that oxidized mitochondrial DNA is tethered to the surface of select mammalian cells through cell surface proteins and heparan sulfate proteoglycans. We demonstrate that cell surface DNA accumulates in large clusters that partially overlap with domains enriched in RNA binding proteins. Finally, we show that human and murine B cell surfaces contain DNA that can be cleared by the secreted nuclease DNASE1L3, and that patients with a *DNASE1L3* missense variant associated with increased risk for autoimmune disease harbor increased levels of surface DNA on B and T cells. Taken together, this work expands the scope of cell surface nucleic acid biology and provides a mechanistic link between cell surface molecules and DNA targeting in autoimmune disease.

## Introduction

Differentiating self versus non-self nucleic acids, especially in the innate immune system, leverages sensors and systems that are topologically contiguous with the extracellular space, including the luminal spaces of the endolysosomal system, thus limiting aberrant immune activation by host intracellular nucleic acids. Still, the extracellular environment is rich in nucleic acid biology, with extracellular, or cell-free (cf) DNA and RNA widely detected in human biofluids, and often serving as biomarkers for disease (reviewed in ^1^). Extracellular RNA is most often comprised of small or fragmented noncoding RNA, thought to arise from the release of apoptotic cellular material ^2–5^. These RNA species are found encapsulated in vesicles ^6^ or bound by RNA binding proteins (RBPs) ^7^, which are thought to serve as protection from secreted nucleases or nucleic acid sensors. More recently, we defined a subset of extracellular RNAs as N-glycosylated and initially characterized them on the cell surface ^8,9^, where the N-glycan serves as a chemical cage to shield underlying RNA modifications that act as Toll-like receptor (TLR) agonists ^10^. GlycoRNAs have also been shown to mediate interactions with endothelial cells ^11,12^, be presented on small extracellular vesicles ^13^, and assume other configurations of glycan-RNA conjugates in exosome vesicles ^14^.

Extracellular DNA has similarly been proposed to result from the release of apoptotic debris ^15^, although the method of nuclease-evasion is often dependent on the source of DNA, with nuclear cfDNA from healthy and cancer patients found in nucleosomes and the majority of cell-free mitochondrial DNA (mtDNA) found in vesicles ^16,17^. Another source of cfDNA comes from neutrophil extracellular traps (NETs), where upon stimulation (often through sensing bacteria), neutrophils release proteins scaffolded on chromatin that can form a web-like structure to trap bacteria ^18^. NETs can also be induced in the context of autoimmune diseases, where stimulation by ribonucleoprotein (RNP) immune complexes leads to the production of NETs that are highly enriched for oxidized mtDNA ^19,20^. Given that oxidized mtDNA is highly inflammatory, either through engagement with the DNA sensor TLR9 ^21^ or the NLRP3 inflammasome ^22^, mtDNA-enriched NETs have also been implicated in autoimmune disease pathology (reviewed in ^22^).

Beyond NETs, nucleic acids have long been linked to autoimmune disease. The first reported autoantibodies from the serum of patients with systemic lupus erythematosus (SLE) were reactive against DNA ^23–25^ and the presence of anti-double-stranded DNA (dsDNA) antibodies is a common clinical diagnostic for SLE ^26^. Moreover, deposits of DNA and anti-double stranded DNA (dsDNA) immune complexes on glomeruli have been linked to lupus nephritis disease progression, suggesting a possible mechanism for DNA-targeting autoantibodies in disease pathology ^27^. Although the molecular mechanism leading to anti-dsDNA production remains unknown, anti-dsDNA antibodies have been found to bind DNA on circulating microparticles (generated from cells undergoing apoptosis), which contain genomic DNA released from apoptotic cells encapsulated in membranes and serve as a possible substrate for recognition by DNA-reactive B cells ^28,29^. Consistent with this idea, the secreted mammalian deoxyribonuclease DNASE1L3 is capable of digesting microparticle-encapsulated DNA ^30,31^ and *Dnase1l3* null mice develop anti-dsDNA antibodies ^31^, suggesting that DNASE1L3 normally clears excess microparticle DNA to prevent the development of autoantibodies directed against *self*-derived dsDNA. In humans, genetic deficiency of *DNASE1L3* results in a monogenic form of dsDNA autoimmune disease with pediatric onset that frequently manifests as SLE or a pre-SLE-like condition ^32–41^.

In our efforts to characterize the nature of the mammalian cell surface as it relates to nucleic acids, we detected surface-localized oxidized mitochondrial DNA that can be released from the plasma membrane upon trypsin treatment. By adapting the use of Assay for Transposase-Accessible Chromatin (ATAC) ^42,43^ to the cell surface, we establish molecular and spatial evidence that cancer cells and primary cells from humans and mice accumulate cell surface DNA. These domains partially overlap with previously characterized cell surface RNA-RBP (csRNP) domains ^44–46^ and are also dependent on heparan sulfate (HS) chains. Finally, we show that the secreted nuclease DNASE1L3 participates in clearing cell surface DNA from primary B and T cells, suggesting a mechanism for regulating the accumulation of mitochondrial DNA on the mammalian cell surface and preventing the development of autoimmune diseases.

## Results

### Mitochondrial DNA can be enzymatically released from the cell surface

Given our previous work demonstrating that RNA is presented on the cell surface in nanoscale clusters containing RBPs and heparan sulfate proteoglycans (HSPGs) ^44,46^, we wondered whether we could selectively release cell surface nucleic acids using trypsin, which has classically been used to cleave cell surface proteins and HSPGs ^47^. Following live cell trypsinization, we extracted nucleic acids from the supernatant (**Figure 1A**). In agreement with recent work reporting cell surface RNA ^8–12,48^, we observed RNase sensitivity of lower molecular weight trypsin-released nucleic acids (**Figure 1B**). Because it has been reported that standard trypsin products contain robust RNase activity ^49^, we also tested cellular release with RNase-free TrypLE. TrypLE-released material phenocopied the trypsin-release profile with particular enhancement of the lower molecular weight products (**Figure 1B**). To our surprise, we also observed DNase-sensitive material across adherent and suspension cancer cell lines (OCI-AML3, A549, and HEK293T cells; **Figure 1B, S1A**), suggesting the possibility that DNA may be present on the cell surface in an analogous manner to RNA. To simplify investigations of DNA-dependent phenotypes, all subsequent enzymatic cell surface releases were performed using trypsin.

**Figure 1.**
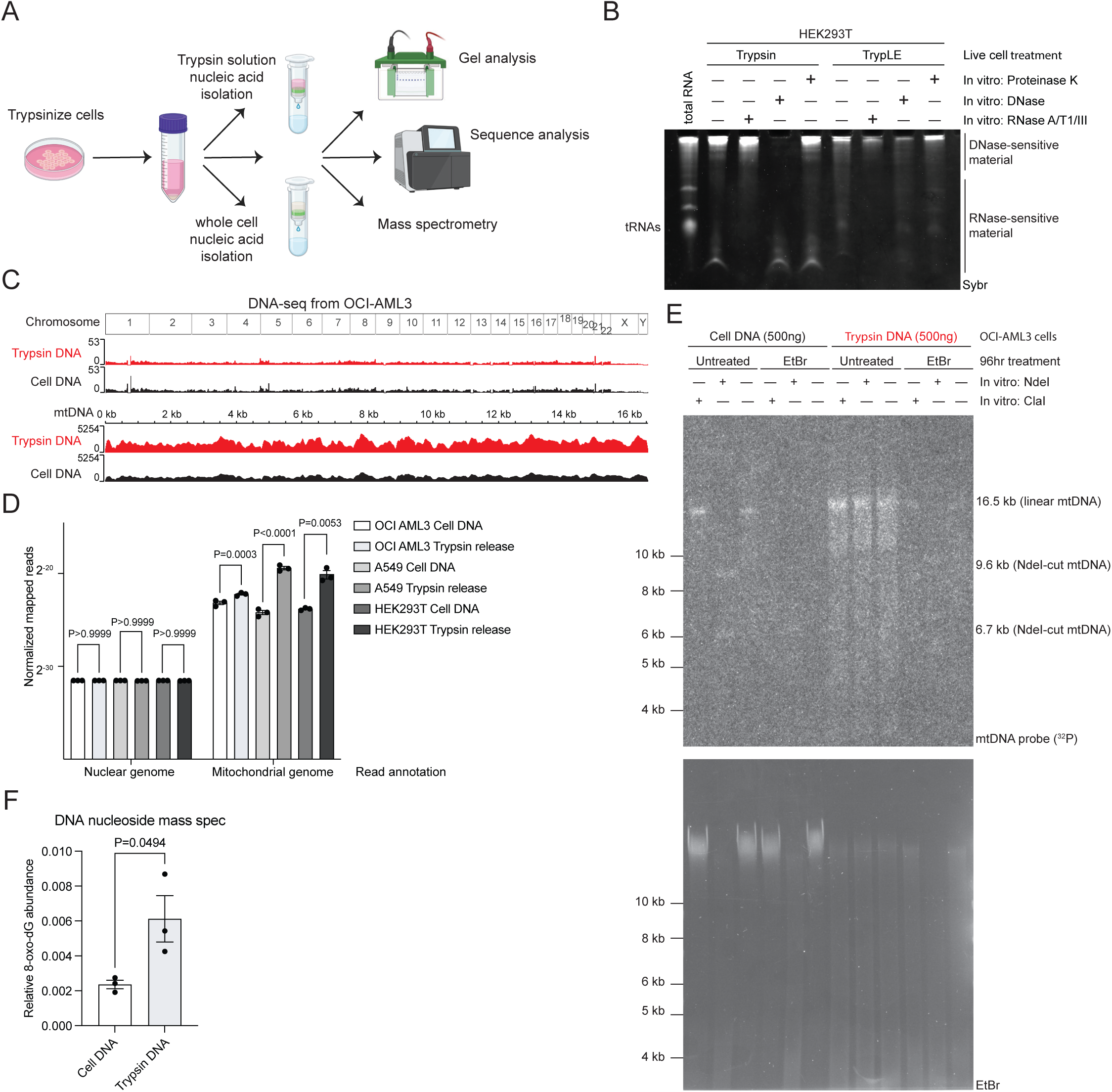
Trypsin releases mitochondrial DNA from cells. A. Schematic of live cell trypsin release, nucleic acid isolation, and downstream analysis. B. Urea PAGE analysis of nucleic acids (SybrGold signal) released from HEK293T cells with trypsin or TrypLE and digested with RNase A/T1/III, TURBO DNase, or Proteinase K. C. Genome browser views of whole cell (Cell DNA) and trypsin-released (Trypsin DNA) DNA from OCI-AML3 cells. Reads were aligned to the nuclear genome (top) and mitochondrial genome (bottom). D. Quantification of trypsin-released DNA from OCI-AML3, A549, and HEK293T cells mapping to the nuclear and mitochondrial genomes. The fraction of reads mapping to each chromosome out of total reads was normalized to chromosome length. *P* values were calculated using a 2-way ANOVA and Tukey’s multiple comparisons test. E. Southern blot analysis of whole cell and trypsin-released OCI-AML3 DNA with and without ethidium bromide-mediated depletion of mitochondrial DNA. DNA was digested with ClaI or NdeI prior to electrophoresis to generate mitochondrial DNA-specific banding patterns. Mitochondrial DNA was detected using a ^32^P-labeled probe against the mitochondrial genome and total DNA was visualized with ethidium bromide prior to membrane transfer. F. Quantification of 8-oxo-2’-deoxyguanosine (8-oxo-dG) relative to 2’-deoxyguanosine in whole cell and trypsin-released DNA from OCI-AML3 cells. *P* value was calculated using an unpaired student’s t-test.

To define the identity of the trypsin-released DNA, we performed fragmentation-based short read DNA sequencing. Across all three cell lines tested, when comparing the whole cell DNA to the trypsin-release samples, we found significant enrichment across the mitochondrial genome (**Figure 1C, D, S1B, S1C, S1D**). We confirmed this by Southern blotting using a labeled probe against human mitochondrial DNA. As expected, mitochondrial signal from whole cell DNA was sensitive to *in vitro* treatment with the mitochondrial DNA-specific restriction enzymes ClaI (linearization of mitochondrial DNA leading to an increase in signal intensity) and NdeI (characteristic fragmentation pattern, ^50^) and was lost upon prolonged treatment with ethidium bromide to deplete mitochondrial DNA ^51^, demonstrating specificity for the selected probes against mitochondrial DNA (**Figure 1E**). We observed an enrichment of mitochondrial DNA signal in trypsin-released DNA, consistent with our DNA sequencing, along with an expected loss of signal upon ethidium bromide treatment, supporting the idea that cell surface mitochondrial DNA released by trypsin originated from cellular DNA, rather than a contaminant from the culture media or trypsin (**Figure 1E**). While trypsin-released DNA was minorly sensitive to *in vitro* ClaI, it was not digested by NdeI (**Figure 1E**), which may be due to the oxidized or fragmented nature of a nucleic acid exposed to the extracellular environment.

Further characterization of the trypsin-released material by mass spectrometry (MS) analysis of modified nucleic acids (SWAMNA, ^52^) confirmed the presence of 8-oxo-2’-deoxyguanosine (8-oxo-dG) (**Figure 1F, S1E**), which is an oxidised form of guanosine commonly found in the mitochondria ^53^. The fractional abundance of 8-oxo-dG was greater in the trypsin-released material, compared to whole cell DNA, consistent with its exposure to an oxidizing environment like the mitochondria. These data suggest that oxidized mitochondrial DNA is present on the surface of mammalian cells and can be liberated by live cell treatment with trypsin.

### *In situ* detection of cell surface DNA reveals clustered organization

To interrogate and establish the features of csDNA *in situ*, we adapted the transposition-based ATAC-Seq/ATAC-See method, which relies on the transposition of dye-conjugated oligonucleotides into accessible regions of dsDNA ^43^. Here, we omitted the initial fixation and permeabilization steps to restrict transposition to cell surface-exposed DNA (**Figure 2A**). Performing cell surface ATAC-See (csATAC-See) with AlexaFluor 647 (AF647)-labeled oligonucleotides on OCI-AML3 leukemic cells revealed puncta clustered on the cell surface (**Figure 2B, Supplementary Movie 1**), but we observe heterogeneous in foci number and intensity across cells. We also observed similarly clustered cell surface puncta on A549 cells, a lung epithelial cancer cell line (**Figure S2A**). csATAC-See signal was dependent on the presence and enzymatic activity of the Tn5 transposase (Tn5) in a time-dependent manner (**Figure 2B, S2B, S2D**) and csATAC-See signal was completely lost upon trypsin treatment, suggesting the ability to label the same population of DNA that can be released with trypsin (**Figure S2C, S2D**). Live cell treatment with benzonase resulted in a near complete loss of cell surface puncta, while live cell RNase treatment had no effect, supporting that DNA is present on the cell surface and accessible to live cell transposition and enzymatic digestion (**Figure 2C**).

**Figure 2.**
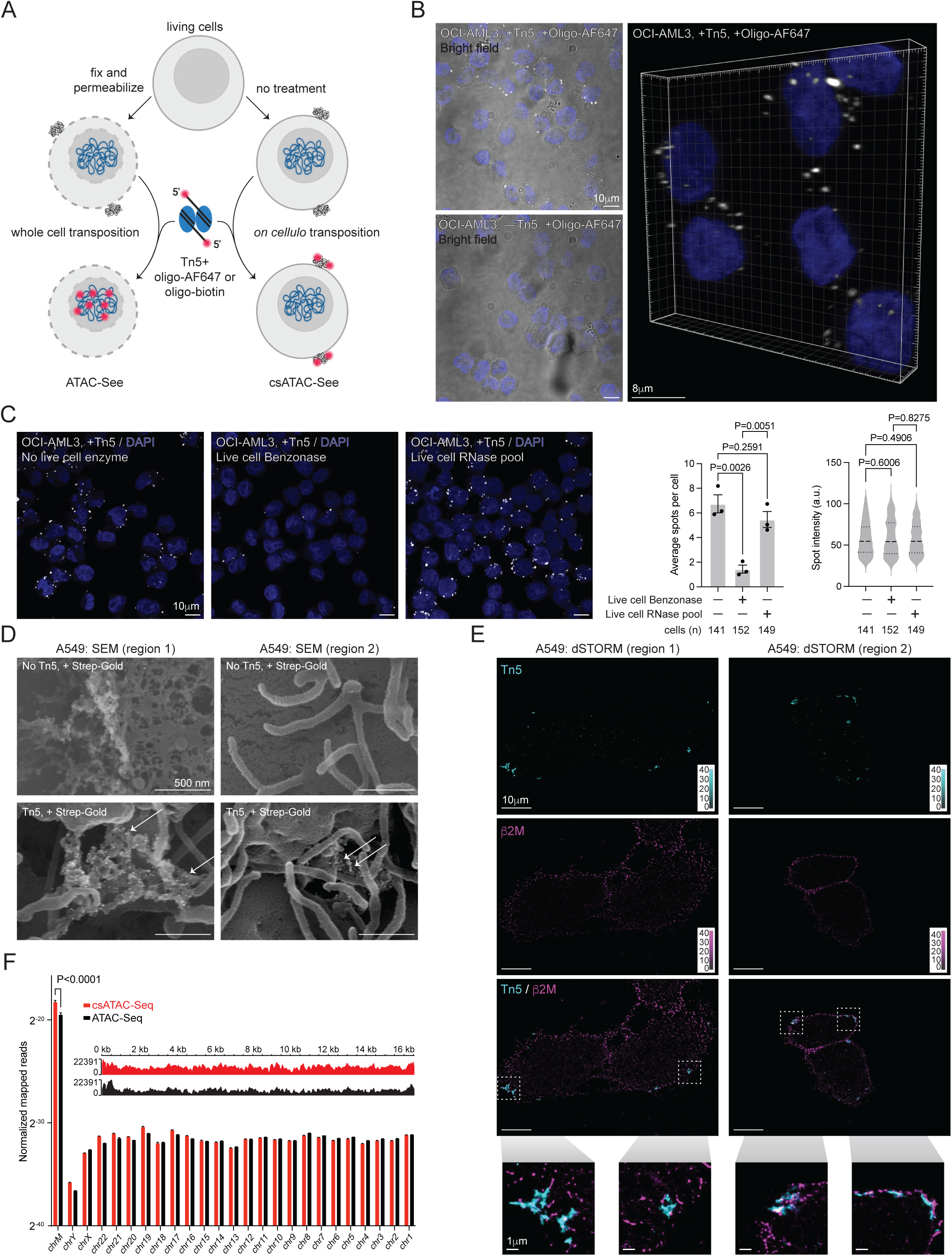
*In situ* detection of cell surface DNA reveals clustered organization. A. Schematic of standard and cell surface (cs) ATAC-See. B. Confocal microscopy of csATAC-See on OCI-AML3 cells with and without Tn5. Oligo-AF647 and DAPI signal from a single z slice are presented with an overlay of a bright-field image. Right: Volumetric reconstruction of 12 z slices. C. Confocal microscopy of csATAC-See on OCI-AML3 cells with and without pre-treatment with benzonase or an RNase cocktail. A single z slice is presented and all z slices were aggregated for quantification of average spot number per cell and spot intensity. The number of cells quantified in each image is shown (n= 3 biological replicates) and *p* values were calculated using an unpaired student’s t-test (average spots per cell) or a Mann-Whitney test (spot intensity). D. Scanning electron microscopy of csATAC-See on A549 cells, visualized using streptavidin-gold (Strep-Gold), with and without Tn5 after a 20 minute labeling reaction. 2 representative regions are shown to depict different cell surface structures. Gold spots are indicated with arrows. E. Super resolution single-molecule localization microscopy of csATAC-See and ꞵ2 microglobulin on A549 cells. The color bar represents the number of detected localizations per 2D histogram bin. Inset: zoomed in regions of csATAC-See spots. F. Quantification of mapped reads per chromosome and a genome browser view of the mitochondrial genome for whole cell and csATAC-Seq from OCI-AML3 cells. *P* values were calculated using a 2-way ANOVA and Tukey’s multiple comparisons test and only *p* values less than 0.05 are presented.

Use of biotinylated csATAC-See oligos allowed us to stain with streptavidin gold (Strep-Gold) and image samples with transmission (TEM) and scanning (SEM) electron microscopy (**Figure 2D, S2E**). We detected gold particles that clearly localized to the external side of the plasma membrane with TEM (**Figure S2E**) and used SEM to further visualize surface-localized gold particles on 2 regions of the cell surface with different surface structures (**Figure 2D**). Our SEM images revealed membrane protrusions which resemble the reported glycocalyx-driven membrane structures ^54^. Next, we performed super resolution single-molecule localization microscopy using direct stochastic optical reconstruction microscopy (dSTORM) on adherent A549 cells to better understand the nature of these cell surface DNA clusters, staining for ꞵ2 microglobulin (ꞵ2M), a component of the major histocompatibility complex I (MHC-I) as a marker of the cell surface (**Figure 2E**) ^44^. Tn5 labeling yielded several irregularly spaced clusters per cell. This is in contrast to ꞵ2M, which appeared more homogenous, as well as our previously reported clusters of csRBPs, which are between 120 and 150 nm in diameter and spaced 250-300 nm apart ^44^.

ATAC-based methods enable sequencing of transposed DNA, allowing an orthogonal method to assess the identity of the csDNA. We prepared standard ATAC-Seq libraries from permeabilized OCI-AML3 cells, as well as csATAC-Seq libraries following live cell transposition and trypsin release prior to library construction, both without post-processing removal of mitochondrial reads. We observed an enrichment of reads mapping to the mitochondrial genome from csATAC-Seq libraries (**Figure 2F**), and decreased peaks at promoters and transcription start sites, relative to standard ATAC-Seq (**Figure S2F, S2G, S2H, Table S3**).

### Cell surface DNA accumulation is controlled by mitochondrial membrane potential, actin polymerization, and heparan sulfate proteoglycans

We next asked which cellular pathways control the presentation of mitochondrial DNA on the cell surface. csATAC-See signal was largely dependent on the presence of mitochondrial DNA, as the number of cell surface puncta were reduced by 60% upon ethidium bromide-mediated depletion of mitochondrial DNA (**Figure 3A**). Short dose ethidium bromide treatment did not lead to a loss of csATAC-See signal, suggesting that it is the loss of mitochondrial DNA itself, rather than the interchelation of ethidium bromide impairing transposition, that contributes to a loss of signal (**Figure S3A**). Similarly, disruption of mitochondrial membrane potential with the mitochondrial membrane uncoupler CCCP also led to a 55% decrease in the number of csATAC-See spots, implicating mitochondrial integrity as another prerequisite for surface accumulation of mitochondrial DNA (**Figure 3B**). csATAC-See signal was also partially dependent on the actin cytoskeleton, as inhibition of actin polymerization with cytochalasin D or latrunculin B each reduced csATAC-See spot number by 40% (**Figure 3C, D, S3B**). Given the ability to release DNA from the cell surface with trypsin, coupled with our previous work demonstrating that csRNPs cluster on the cell surface in a heparan sulfate-dependent manner ^46^, we also tested csATAC-See signal sensitivity to live cell addition of a heparinase cocktail (HSase). csATAC-See spots diminished 60% following heparinase treatment, suggesting a potentially common mechanism for heparan sulfate proteoglycans in tethering nucleic acids to the cell surface (**Figure 3E**). We also evaluated dependence on nanotubes (**Figure S3C**), endoplasmic reticulum (ER) to Golgi transport (**Figure S3D**), and mitochondrial reactive oxygen species (ROS) (**Figure S3E**), finding no significant effects with the inhibitors used. It is notable that inhibition of nanotubes does not affect csATAC-See signal, as recent studies implicated tunneling nanotubes (TNT) in the transport of intact mitochondria between cells ^55,56^, arguing against such a mechanism for the presentation of mitochondrial DNA on the surface.

**Figure 3.**
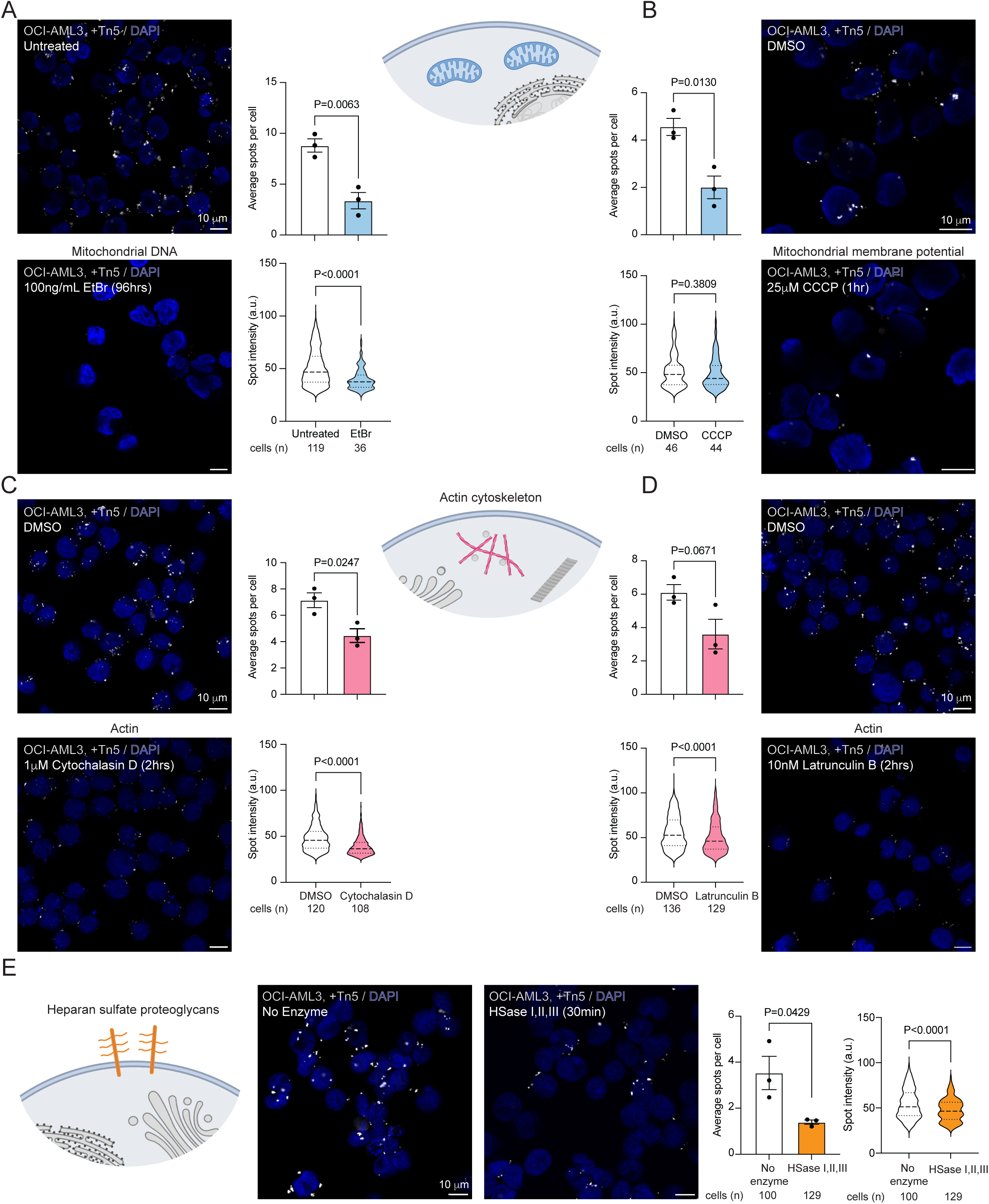
Cell surface DNA accumulation is dependent on mitochondrial integrity, actin polymerization, and heparan sulfate proteoglycans. A. Confocal microscopy of csATAC-Seq on OCI-AML3 cells with and without ethidium bromide-mediated mitochondrial DNA depletion. Treatment time and conditions are indicated, along with the number of cells quantified in each image (n= 3 biological replicates). *P* values were calculated using an unpaired student’s t-test (average spots per cell) or a Mann-Whitney test (spot intensity). B. As in A, treated with the mitochondrial membrane potential uncoupler Carbonyl Cyanide m-Chlorophenylhydrazone (CCCP). C. As in A, treated with the actin inhibitor Cytochalasin D. D. As in A, treated with the actin inhibitor Latrunculin B. E. As in A, treated with a live cell heparinase cocktail (HSase I, II, III).

### Cell surface DNA localizes near RNA binding proteins and integrins

Loss of csATAC-See signal after HSase suggested certain membrane proteins, like HSPGs, play a role in the cell surface tethering of DNA. To define the local proteome of these DNA species, we used biotinylated oligonucleotides in the csATAC protocol, followed by staining with streptavidin coupled to horse radish peroxidase (HRP). Subsequent proximity labeling with biotin phenol was used to define the DNA-proximal proteome on OCI-AML3 cells (csATAC-protein proximity labeling, csATAC-pPL) ^44,57^. This yielded 195 significant hits with an enrichment of more than 2-fold compared to a parallel csATAC-pPL reaction omitting Tn5 (**Figure 4A, S4A**). Gene ontology analysis revealed shared functions in RNA binding and integrin binding, among others, as well as proteins classically localized to the cell surface (**Figure 4B**). Given the number of hits with functions relating to RNA binding, we intersected the list of Tn5 proximal proteins with proximity labeling datasets for the cell surface RBPs DDX21, nucleolin (NCL) ^44^, and nucleophosmin (NPM1) ^45^ (**Figure 4C, S4B**). We detected 100 hits shared between Tn5 and at least one cell surface RBP, 36 of which were annotated RBPs (**Figure 4C, S4C**).

**Figure 4.**
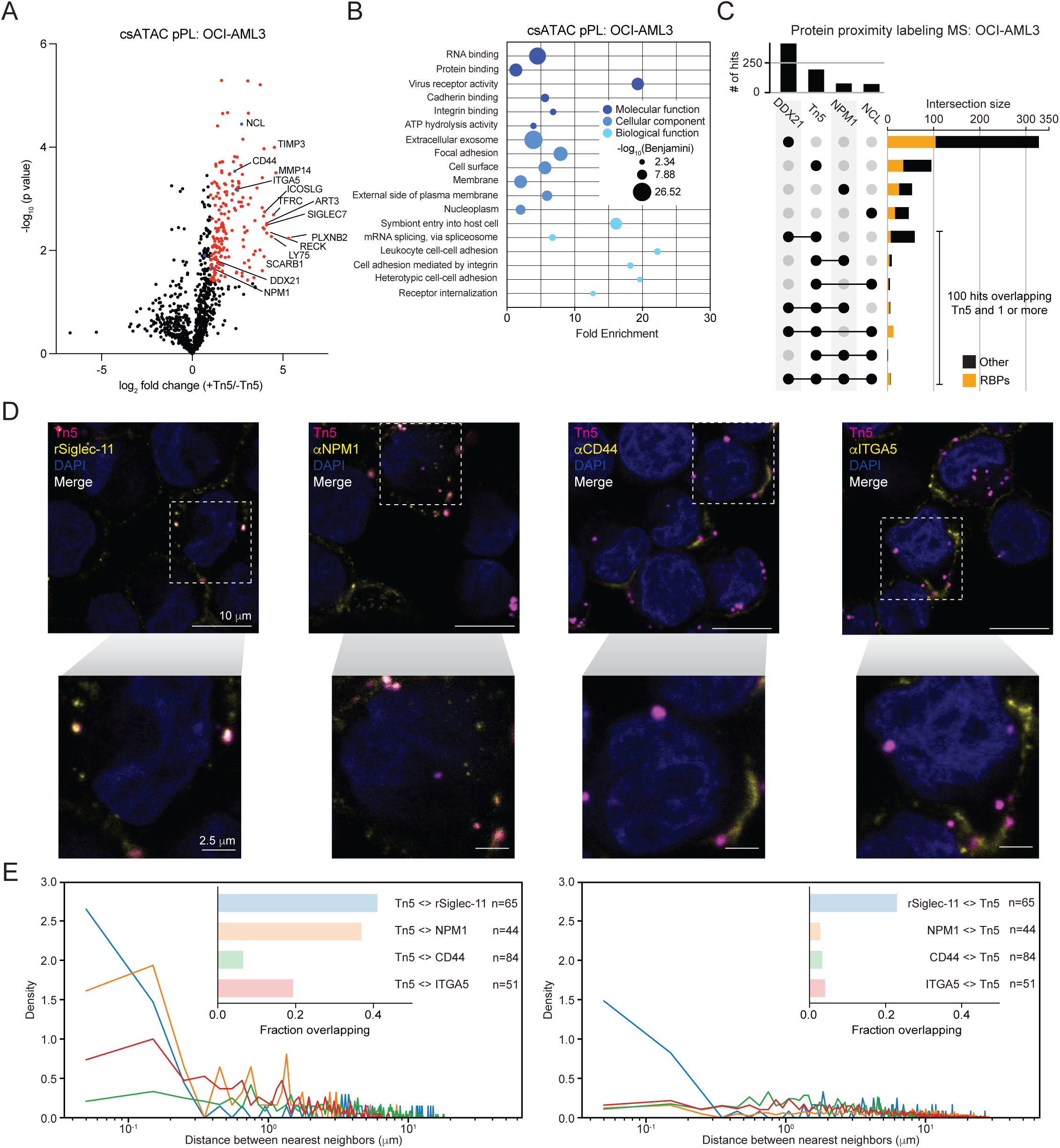
Molecular organization and presentation of cell surface DNA. A. Volcano plot of csATAC protein proximity labeling (pPL) mass spectrometry (MS) on OCI-AML3 cells (n= 4 biological replicates). Proteins enriched more than 2-fold over the no Tn5 condition and with a significant *p* value determined by a Welch’s t-test are colored in red. Proteins of interest, including those for which we have previously generated proximity labeling datasets ^44,45^ are indicated in blue and the identities of the top 10 enriched hits are indicated. B. Gene ontology (GO) analysis of molecular function, cellular component, and biological function for proteins identified by csATAC pPL MS. C. Intersection analysis of csATAC pPL with cell surface RNA binding protein (csRBP) pPL datasets from OCI-AML3 cells ^44,45^. The fraction of RBP hits is indicated in yellow. D. Confocal microscopy of csATAC-See co-localization with Siglec-11, NPM1, CD44, and ITGA5 on OCI-AML3 cells. A merged image representing a single z slice is shown. E. Nearest neighbor distance analysis of the indicated reagent or antibody signal with csATAC-See signal (antibody near Tn5 spots). Average pair densities of a single z slice from 3 ROIs were plotted in a density histogram. A bar plot of a Manders coefficient representing the fraction of overlapping spots for each csATAC-See-antibody pair is shown. F. Same as E, for csATAC-See signal with the indicated reagent or antibody signal (Tn5 near antibody spots).

Given the overlap between csDNA- and csRBP-proximal proteins, we performed csATAC-See followed by cell surface antibody labeling and confocal microscopy to directly visualize the spatial distribution of DNA and cell surface RBPs, as well as other enriched hits (**Figure 4D, S4D**). We also assessed the co-localization between cell surface DNA and RNA by staining cells with recombinant Siglec-11, which we have shown binds the cell surface in an RNA-dependent manner ^8,46^ and binds in proximity of other cell surface RBPs ^46^. Quantification of co-localization revealed a high degree of co-localization between cell surface DNA and antibody signal, particularly for Siglec-11 and NPM1, with most Tn5-labeled spots overlapping with Siglec-11 and NPM1 spots (**Figure 4E**), although since Siglec-11 and NPM1 spots are more abundant, not all Siglec-11 and NPM1 spots overlapped with cell surface DNA (**Figure 4E**, right panel). This suggests that DNA and RNA occupy partially overlapping domains on the cell surface. csDNA spots are also present near the cell surface glycoprotein CD44 and integrin subunit alpha 5 (ITGA5), but with less overlap, in agreement with our proximity labeling indicating shared molecular neighborhoods.

### DNA is present on the surface of mammalian B cells and monocytes

Since our characterization of csDNA was conducted on immortalized cell lines, we wondered whether DNA also accumulates on the surface of healthy cells. Given that OCI-AML3 are myeloid of origin, we started with immune cells. We first validated flow cytometry as a new readout for csATAC-See by benchmarking our staining on OCI-AML3 cells and observed robust AF647 signal on live cells with Tn5, compared to a negative control that only contained AF647-labeled oligonucleotides (**Figure 5A, S5A**). Additionally high AF647 staining was abolished when the transposition reaction was performed on ice, recapitulating our csATAC-See confocal data (**Figure 5A**). To determine whether csATAC-See signal was associated with apoptosis, we stained live cells with Annexin V following csATAC-See and found that Annexin V-cells stained positive for csATAC-See, indicating that live, non-apoptotic cells display csDNA (**Figure 5B, S5B, S5C**).

**Figure 5.**
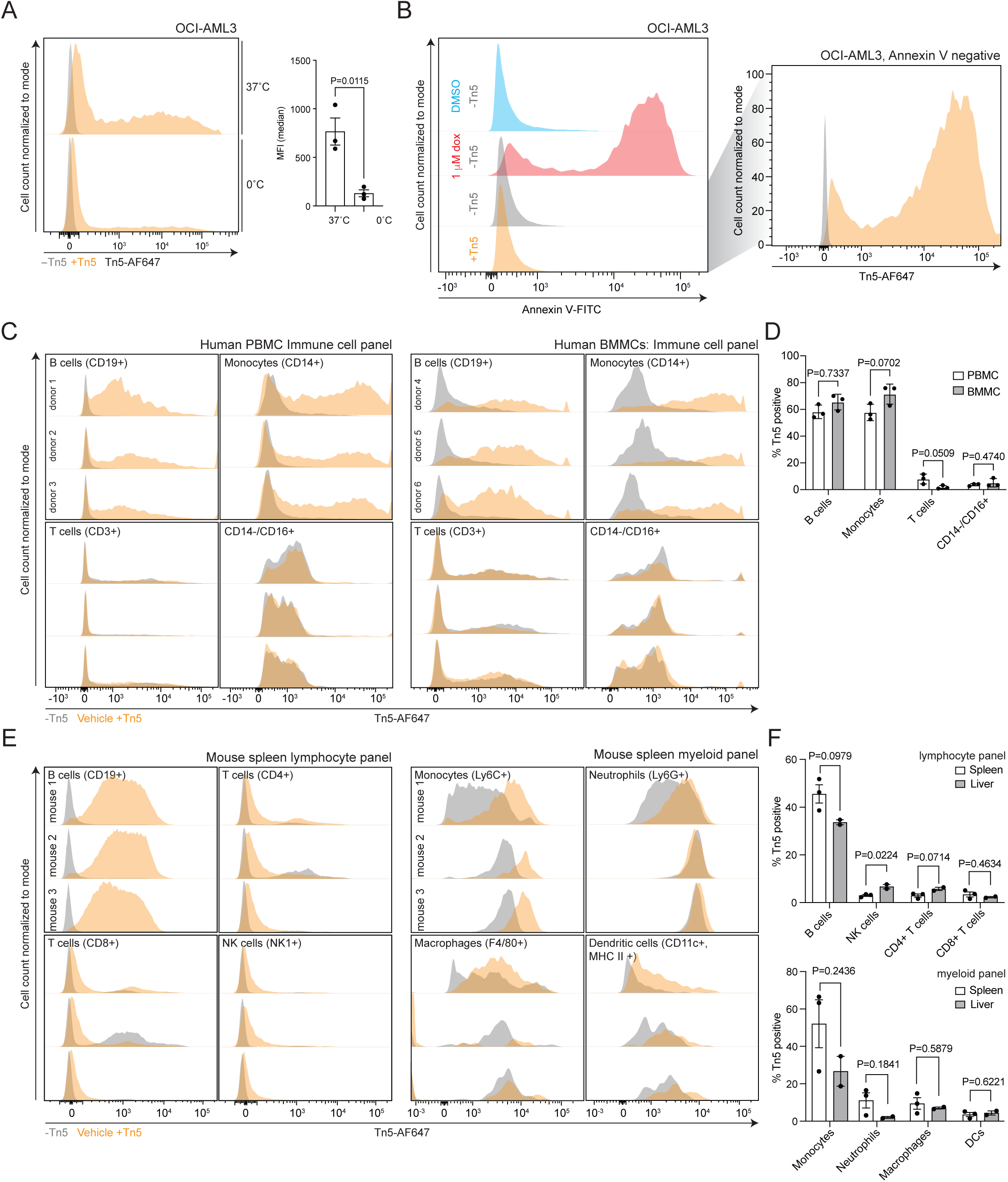
Mammalian B cells and monocytes display cell surface DNA. A. Live cell flow cytometry analysis of OCI-AML3 cells stained with AF647-labeled oligos, with (orange histograms) or without (gray histograms) Tn5 at 37°C or on ice. Right: Quantification of csATAC-See signal. Median fluorescence intensity was calculated by subtracting the median on cells without Tn5. *P* values were calculated using an unpaired student’s t-test (n= 3 biological replicates). B. Annexin-V staining on OCI-AML3 cells treated for 24 hours with DMSO or 1 μM doxorubicin, with or without prior cs-ATAC-See staining. Right: csATAC-See signal on Annexin V negative OCI-AML3 cells. C. Live cell flow cytometry analysis of human peripheral blood mononuclear cells (PBMCs) and human bone marrow mononuclear cells (BMMCs) stained with AF647-labeled oligos and without Tn5 (gray histograms) or with Tn5 (orange histograms). Results from 3 donors for each sample type are displayed. D. Quantification of PBMCs and BMMCs gated as Tn5 positive for each cell type. *P* values were calculated using an unpaired student’s t-test (n= 3 donors for each sample type). E. Live cell flow cytometry of mouse spleen cells using a lymphocyte and myeloid antibody panel (n= 3 mice). F. Quantification of spleen and liver cells gated as Tn5 positive for each cell type. *P* values were calculated using an unpaired student’s t-test (n= 3 mice for spleen, 2 mice for liver).

We next performed csATAC-See on healthy human peripheral blood mononuclear cells (PBMCs) and human bone marrow mononuclear cells (BMMCs) to assess csDNA levels by flow cytometry. Both PBMCs and BMMCs also showed positive staining relative to the negative control across 3 different donors each (**Figure 5C, 5D, S5C, S5D, S5F, S5G**), indicating that csDNA is not limited to immortalized cells in culture. On PBMCs and BMMCs, B cells and monocytes had high levels of specific Tn5 staining, while T cells and CD14-/CD16+ cells (which include NK cells) did not show staining above background (**Figure 5C, 5D**). We noted some variability among donors, which may reflect slight differences in cell type composition (**Figure S5D, S5H**) or biological differences across individuals.

To determine if csDNA accumulation was a phenomenon limited to human cells, we harvested leukocytes from mouse spleens and livers and performed csATAC-See with antibody panels capturing an expanded set of lymphocyte and myeloid cell types (**Figure 5E, 5F, S6, S7**). Similar to what we observed on human PBMCs and BMMCs, B cells harvested from the spleen and liver displayed robust csATAC-See signal over background levels, while T cells and NK cells showed lower or more variable staining above the no enzyme control (**Figure 5E, 5F, S6A**). On myeloid cells, we detected high signal-to-noise of csATAC-See signal on monocytes in the spleen, but less robust in the liver and little to no csATAC-See signal on neutrophils, dendritic cells (DCs), or macrophages in either organ (**Figure 5E, 5F, S6A, S7**). Taken together, this suggests that csDNA accumulation is conserved among humans and mice and, likewise, that B cells and monocytes are the primary sites of csDNA accumulation in the healthy immune system.

### DNASE1L3 removes cell surface DNA from B cells

The presence of csDNA on healthy B cells and monocytes, but not other tested immune cell types, was intriguing, as such cell type specificity could hint at either a functional relevance for csDNA or regulated mechanisms controlling the accumulation of DNA across different cell types. The extracellular space is normally considered to be inhospitable for nucleic acids, especially RNA with many secreted RNases (reviewed in ^58^). However, mammals also secrete DNases into the extracellular environment, including DNASE1, which is primarily secreted by cells in the digestive tract ^59^, and DNASE1L3, which is secreted by dendritic cells, macrophages, and to a lesser extent, B cells, in the liver and spleen ^31,59^. Since DNASE1L3 has been reported to have activity on protein- or lipid-complexed DNA (in the context of microparticles in the extracellular space) and loss of DNASE1L3 functionality has been associated with increased antibodies against double-stranded DNA (dsDNA, ^31^) and autoimmune disease ^32^, we wondered whether csDNA might be an endogenous target of DNASE1L3. To test this, we performed csATAC-See on human PBMCs pre-treated with recombinant wild type DNASE1L3 or a vehicle (the DNASE1L3 R206C variant, which causes a defect in protein secretion and is not detectable when recombinantly expressed, nor displays enzymatic activity on isolated nuclei or genomic DNA ^60^) (**Figure 6A, 6B, S6F, S6G**). Recombinant DNASE1L3 reproducibly decreased csATAC-See signal on B cells, although the effect of DNASE1L3 on monocyte csATAC-See signal was variable across donors (**Figure 6A, 6B**). We observed the same trend on BMMCs treated with recombinant DNASE1L3 (i.e. a decrease in cell surface DNA on B cells, but not monocytes), although the trend on B cells was weaker and more variable across donors, compared to PBMCs (**Figure 6C, 6D, S6G**).

**Figure 6.**
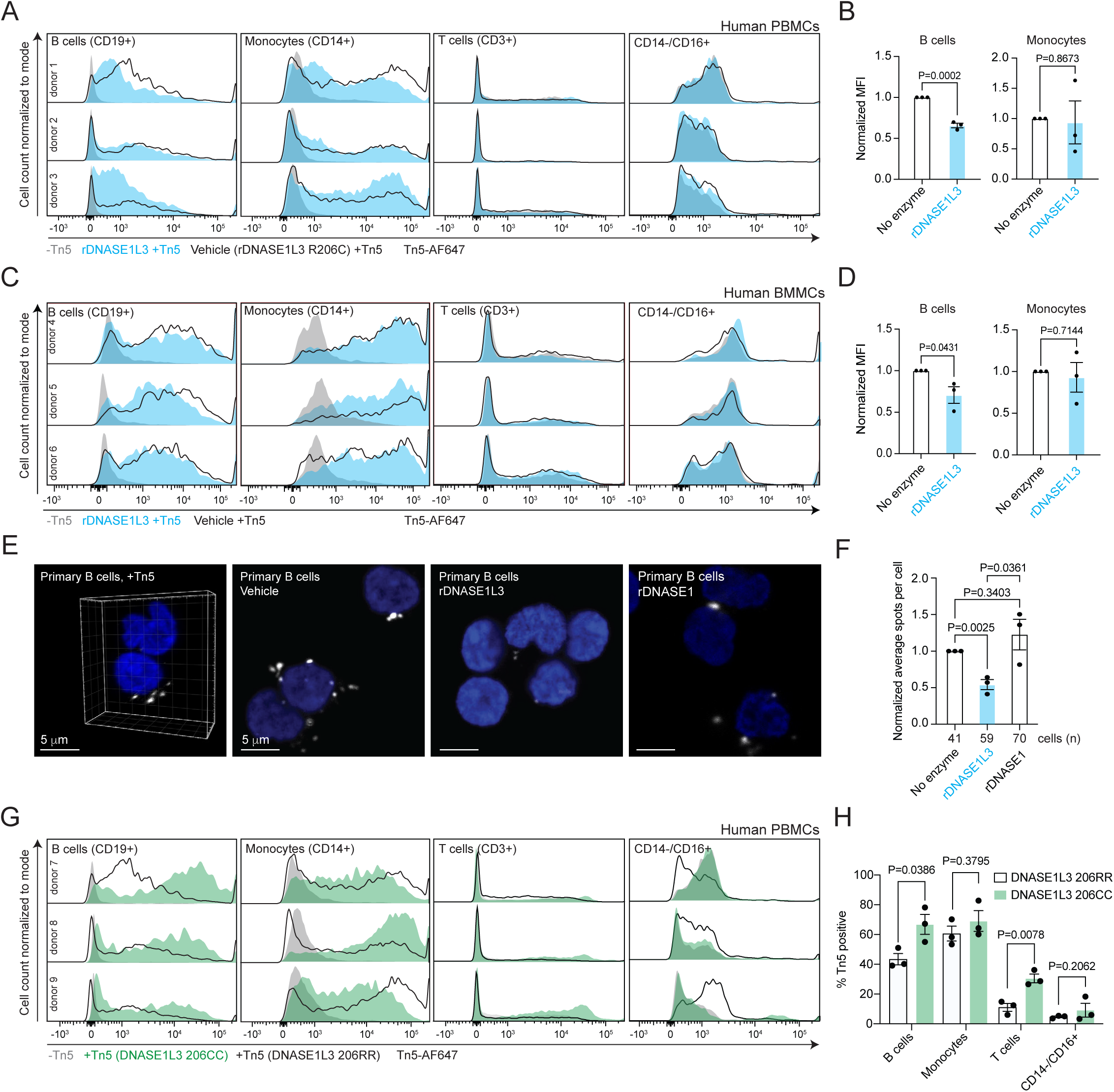
DNASE1L3 surveils DNA on the surface of B cells. A. Live cell flow cytometry analysis of human PBMCs stained with AF647-labeled oligos and Tn5 with recombinant DNASE1L3 (blue histograms) or without Tn5 (gray histograms). Black-outlined histograms represent the same samples treated with a vehicle control (DNASE1L3 R206C). Results from 3 donors are displayed for each immune cell type. B. Normalized mean fluorescence intensity (MFI, geometric mean) was calculated by subtracting the MFI on cells without Tn5, then further normalized to cells without recombinant DNASE1L3. *P* values were calculated using an unpaired student’s t-test (n= 3 donors). C. Live cell flow cytometry analysis of human bone marrow mononuclear cells (BMMCs) stained with AF647-labeled oligos and Tn5 with recombinant DNASE1L3 (blue histograms) or without Tn5 (gray histograms). Black-outlined histograms represent the same samples treated with a vehicle control (DNASE1L3 R206C). Results from 3 donors are displayed for each immune cell type. D. Normalized mean fluorescence intensity (MFI, geometric mean) on BMMCs, as in B. *P* values were calculated using an unpaired student’s t-test (n= 3 donors). E. *In vitro* nuclei and genomic DNA digestion with recombinant DNASE1L3 (wild type and R206C/vehicle) or DNASE1. F. Quantification of PBMCs and BMMCs gated as Tn5 positive for each cell type, with or without pre-treatment with recombinant DNASE1L3. *P* values were calculated using an unpaired student’s t-test (n= 3 donors). G. Volumetric reconstruction (from 12 z slices) of csATAC-See on primary isolated B cells and representative single z slices of csATAC-See on primary isolated B cells with and without pre-treatment with recombinant DNASE1L3 or DNASE1. H. Quantification of average csATAC-See spots per cell after recombinant DNASE1L3 or DNASE1 treatment, normalized to untreated cells. The number of cells quantified is shown and *p* values were calculated using an unpaired student’s t-test (n= 3 donors). I. Live cell flow cytometry analysis of human peripheral blood mononuclear cells (PBMCs) stained with AF647-labeled oligos and with (DNASE1L3 206RR/wild type, black histograms and DNASE1L3 206CC/homozygous risk allele, green) or without (gray histograms) Tn5. Results from 3 donors per genotype are displayed for each immune cell type. J. Quantification of PBMCs gated as Tn5 positive for each cell type, for wild type and homozygous mutant *DNASE1L3* patients. *P* values were calculated using an unpaired student’s t-test (n= 3 donors).

To visualize the DNASE1L3 B cell phenotype, we isolated B cells from fresh PBMCs, treated cells with recombinant DNASE1 or DNASE1L3, and monitored csATAC-See signal by confocal microscopy (**Figure 5E**). DNASE1L3, but not DNASE1, partially and reproducibly removed cell surface DNA (**Figure 5E, 5F**). Finally, to assess if this activity could occur in humans, we examined PBMCs from patients homozygous for the rs35677470 risk allele, which encodes the R206C substitution in DNASE1L3 and is associated with increased risk for the development of SLE and scleroderma ^60–63^. It has been reported that patients with homozygous mutations secrete 10-fold less DNASE1L3 than patients with the wild type allele ^60^. In individuals with this variant, we detected an increase in csATAC-See signal on B cells, as well as increased signal on T cells to levels above background (**Figure 5G, 5H**). Together, these results support a role for DNASE1L3 in maintaining homeostatic levels of csDNA on B and suggesting additional *in vivo* targets that include T cells.

## Discussion

The cell surface has rapidly emerged as a novel environment for studying nucleic acid biology. Here we present evidence that mitochondrial DNA is stably associated with the cell membrane, anchored in part by heparan sulfate proteoglycans in a manner analogous to the tethering of RNA and RBPs on the cell surface ^46^. We also show that csDNA is not limited to cultured immortalized cells, as it is also on healthy primary cells from humans and mice, where it displays specificity for B cells and monocytes. Given the ability for DNASE1L3 to liberate csDNA from primary B cells, we propose that the cell surface may act as a reservoir to store immunogenic nucleic acid ligands such as oxidized mitochondrial DNA, where DNASE1L3 acts as a homeostatic mechanism to metabolize csDNA and limit its accumulation to levels that can be tolerated by the immune system. DNASE1L3 has previously been reported to act on cfDNA or apoptotic DNA presented on microparticles ^31,64^, the latter of which can serve as a target of autoantibodies from SLE patients ^29,65^. Notably, the positively charged C-terminal region of DNASE1L3, which is critical for its activity on membrane-complexed DNA *in vitro*, has been proposed to act as a cell-penetrating peptide that can facilitate membrane binding in the context of microparticles ^31^. Our data describing a function for DNASE1L3 in digesting cell surface-bound DNA suggests that its ability to target membranes through the C-terminal domain may facilitate its interaction with the cell surface, thereby expanding the scope of DNASE1L3 targets beyond microparticle DNA and cfDNA.

Clinically, DNASE1L3 has best been characterized for its role in autoimmunity by preventing the accumulation of anti-dsDNA autoantibody targets. DNASE1L3 KO mice develop autoantibodies against dsDNA and features of SLE, including IgG deposition in kidney glomeruli and glomerulonephritis ^31^. Similarly, SLE patients with loss-of-function mutations in the *DNASE1L3* gene also develop anti-dsDNA antibodies ^32^, indicating a link between effective clearance of DNA in the extracellular environment and the generation of anti-dsDNA antibodies. Although the exact mechanism of anti-dsDNA autoantibody generation remains unknown, as is the source of DNA that initiates antibody production, our data indicating that csDNA is accessible to and a substrate of DNASE1L3 suggests the possibility that csDNA, like microparticle DNA, may be recognized by patient autoantibodies, or lead to the production of autoantibodies under conditions where there are abnormally high levels of csDNA, such as mutations in the *DNASE1L3* gene.

Our results also reveal several biological pathways implicated in the presentation and accumulation of csDNA. csDNA presentation is at least partly dependent on the actin cytoskeleton, which may reflect vesicle-mediated transport to the surface, or cytoskeletal anchoring of DNA present on the surface or in the extracellular environment. We also show that csDNA presentation is highly sensitive to the digestion of HS chains, which is consistent with our previous work demonstrating that HS chains anchor csRNPs ^46^, and suggests a common mechanism for tethering nucleic acids to the cell surface. Further, the spatial overlap of csATAC-See signal with csRBP clusters points towards common domains where DNA and RNA are present. Since live cell RNase digestion did not impact csATAC-See signal, this suggests that although csDNA and RNA may co-localize, csDNA accumulation is not dependent on intact cell surface RNA. Still, it remains unknown whether there are any functional interactions between DNA and RNP clusters on the cell surface, especially given that many of the csRBPs we have previously characterized have also been reported to bind DNA ^44,45,66,67^. Beyond HS-mediated tethering, we also show that csDNA presentation is dependent on various aspects of mitochondrial biology, consistent with the enrichment of mitochondrial DNA we detect on the surface. It is interesting that csDNA presentation relies on healthy mitochondria with functional oxidative phosphorylation, as CCCP-mediated membrane uncoupling has been reported to lead to a loss of mitochondrial DNA secretion by neutrophils ^20^, suggesting that the cell surface mitochondrial DNA presentation we describe here and mitochondrial DNA extrusion in the context of neutrophil extracellular traps (NETs) rely on different cellular contexts.

Our results demonstrating that mtDNA accumulates on the surface of mammalian cancer cells and immune cells is reminiscent of recent reports that human and murine red blood cells bind CpG DNA, including exogenous mtDNA, pathogen DNA, and mtDNA that accumulates during inflammation and after cell death, in a manner dependent on surface presentation of TLR9 ^68,69^. This is thought to be a mechanism to clear red blood cells in response to pathogens or inflammatory stress, as DNA binding through surface-expressed TLR9 leads to accelerated erythrophagocytosis. It is unclear whether the DNA scavenged by red blood cells is stably tethered on the cell surface or internalized from the extracellular environment prior to clearance of CpG DNA-containing red blood cells. Similarly, platelets have also been reported to act as cfDNA scavengers that bind and internalize cfDNA through clathrin-mediated endocytosis of extracellular vesicles or direct uptake of membrane-free DNA ^70^. Platelet-scavenged cfDNA primarily maps to the nuclear genome and its internalization does not rely on the actin cytoskeleton ^70^, in contrast to what we have characterized for csDNA. Still, these studies and ours point towards the importance of dedicated mechanisms to limit circulating cfDNA and csDNA accumulation, which we hypothesize is critical to prevent immune responses under homeostatic conditions.

While we have characterized the identity of csDNA and demonstrated its conservation across primary and immortalized mammalian cells, this work does not address the source of csDNA— whether csDNA is presented on the same cell of origin or if secreted or cfDNA can accumulate on the surface of any cell, regardless of origin. Following this, although our flow cytometry results indicate that csDNA is present on live cells, we are also unable to address whether csDNA originated from live or apoptotic cells, as has been suggested for microparticle DNA. Additionally, it remains unclear whether csDNA is functional, or simply the passive result of a cell existing in the nucleic acid-rich extracellular space. Further work leveraging the tools herein could reveal answers to these questions.

Finally, in this work we also describe the development of csATAC-See, a new variant of ATAC-Seq/See that is specific for the cell surface. We show that csATAC-See can be applied to cell culture models and primary cells, displays sensitivity to live cell nuclease treatment, and is amenable to spatial multi-omics profiling including next-generation DNA sequencing, confocal, super resolution, or electron microscopy, and the identification of proximal proteins by mass spectrometry. Similar methods have recently been developed for RNA ^71,72^ and chromatin ^73^, although these methods rely on prior knowledge of the sequence of interest and chemical fixation, which may disrupt organization of the cell surface ^74^. Thus, csATAC-See represents a rapid method for interrogating csDNA from a wide variety of cell or sample types without the need for additional design considerations. We anticipate that csATAC will be a useful tool in continuing to probe csDNA in health and disease.

## Figure Legends

**Figure S1.**
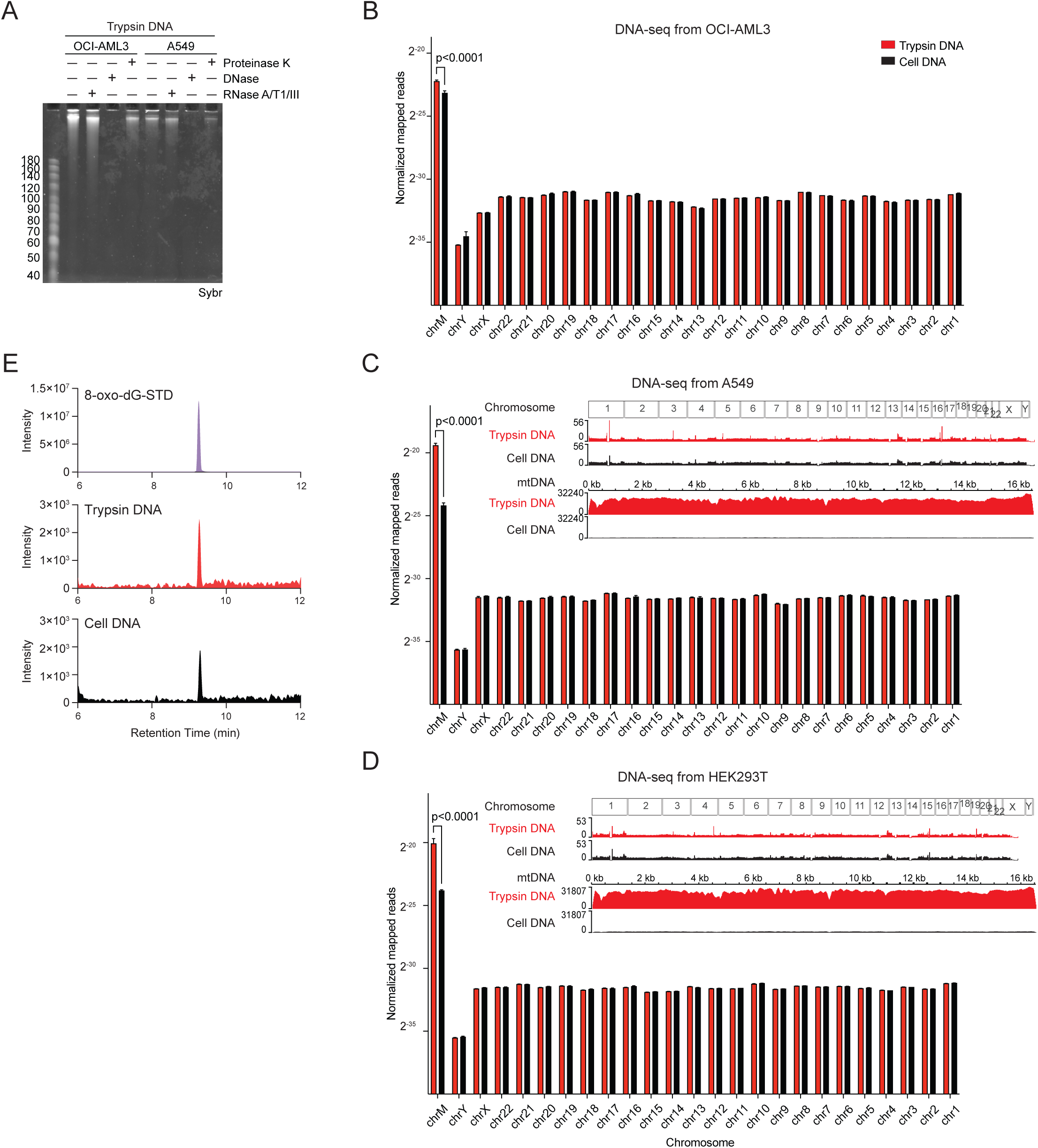
Trypsin releases mitochondrial DNA from immortalized adherent and suspension cells. A. Urea PAGE analysis of nucleic acids (SybrGold signal) released from OCI-AML3 and A549 cells with trypsin and digested with RNase A/T1/III, TURBO DNase, or Proteinase K. B. Quantification of mapped reads per chromosome for whole cell and trypsin-released DNA from OCI-AML3 cells. *P* values were calculated using a 2 way ANOVA and Tukey’s multiple comparisons test and *p* values less than 0.05 are presented. C. Genome browser views and quantification of mapped reads per chromosome for whole cell and trypsin-released DNA from A549 cells, with statistics described in B. D. Same as C, for DNA from HEK293T cells. E. MS2 spectra of an 8-oxo-dG standard and representative trypsin and cell DNA samples.

**Figure S2.**
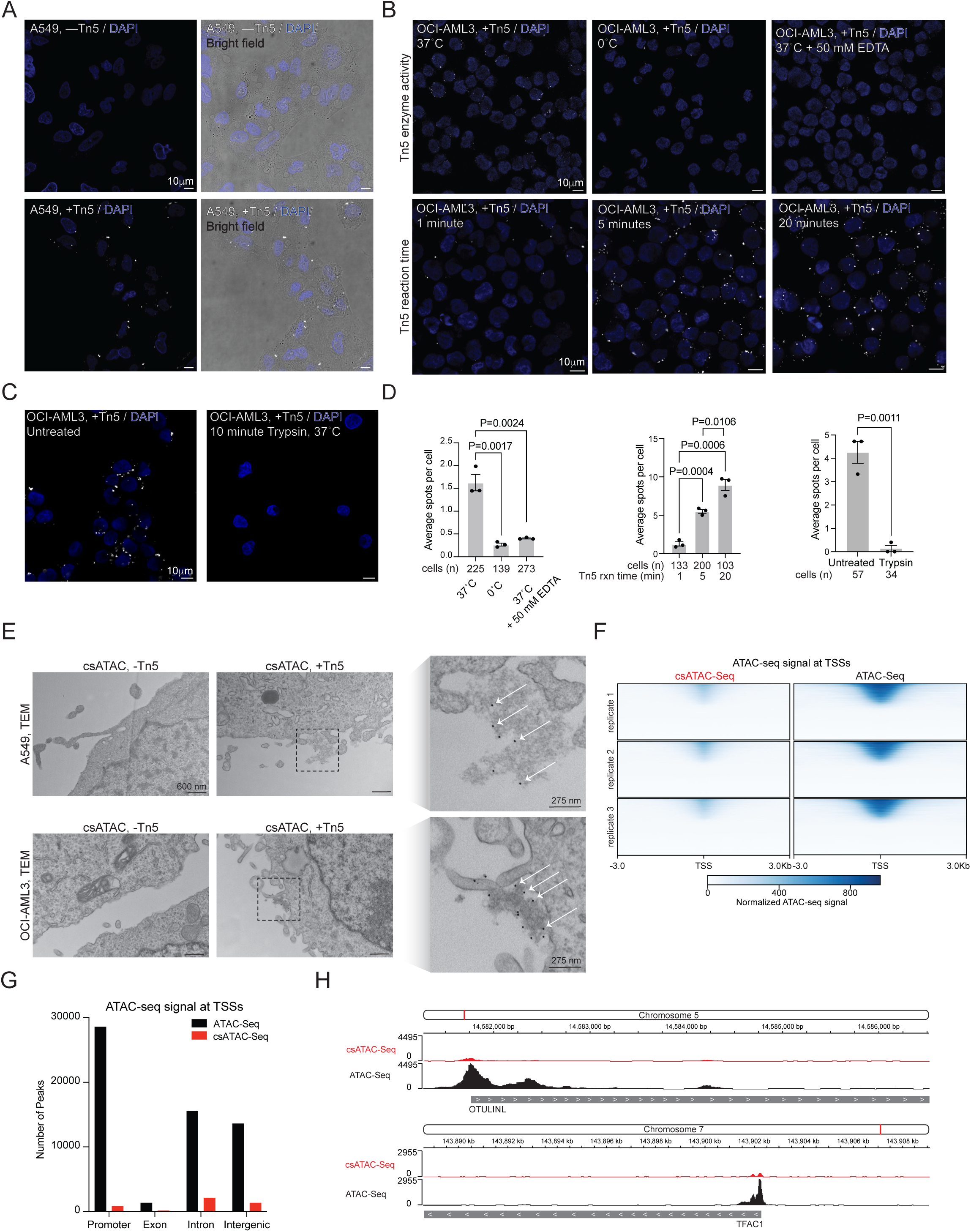
Cell surface ATAC-See labels DNA in a time- and enzyme-dependent manner. A. Confocal microscopy of csATAC-See on A549 cells with and without Tn5. A single representative z slice is shown. B. Top: Confocal microscopy of csATAC-See on OCI-AML3 cells performed on ice, or with the addition of EDTA to inactivate Tn5 enzymatic activity. Bottom: Confocal microscopy of csATAC-See on OCI-AML3 cells with different labeling reaction times. C. Confocal microscopy of csATAC-See on OCI-AML3 cells with or without pre-treatment with trypsin. D. Quantification of csATAC-See signal in B and C. The number of cells quantified in each image is shown (n= 3 ROIs) and *p* values were calculated using an unpaired student’s t-test. E. Transmission electron microscopy of csATAC-See on OCI-AML3 and A549 cells, visualized using streptavidin-gold (Strep-Gold) with and without Tn5 after a 5 minute labeling reaction. Gold spots are indicated with arrows on the zoomed in images. F. Heat map of standard and csATAC-Seq signal at transcription start sites (TSS) for 3 biological replicates, with color intensity representing normalized ATAC-Seq signal. G. Number of peaks for genomic features detected for standard and csATAC-Seq. Peaks were called from 3 biological replicates using MACS2. H. Genome browser views of standard and csATAC-Seq signals at representative promoter regions.

**Figure S3.**
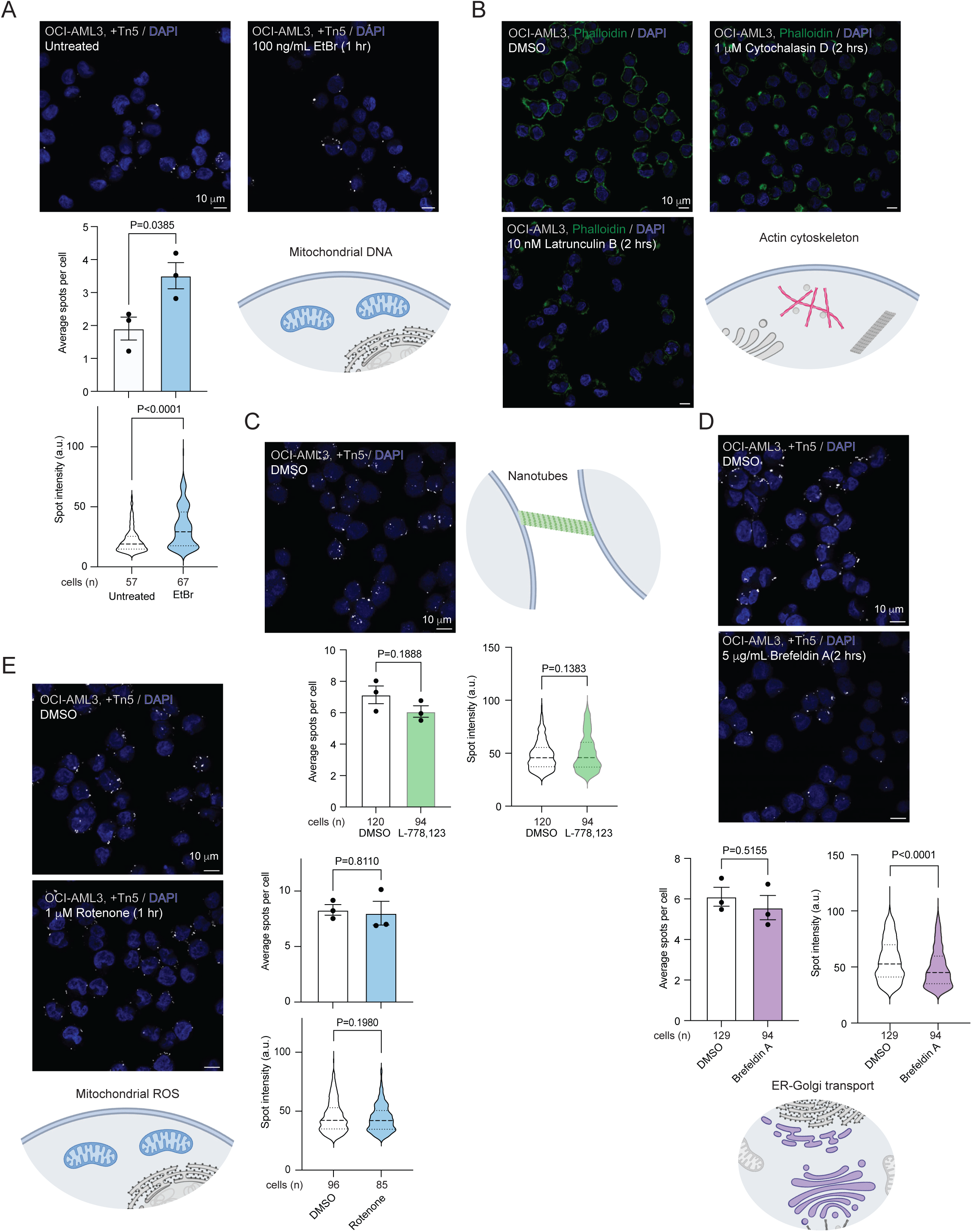
Mechanisms of cell surface DNA presentation. A. Confocal microscopy of csATAC-Seq on OCI-AML3 cells with and without 1 hour ethidium bromide treatment. Treatment time and conditions are indicated, along with the number of cells quantified in each image (n= 3 biological replicates). *P* values were calculated using an unpaired student’s t-test (average spots per cell) or a Mann-Whitney test (spot intensity). B. Phalloidin staining of actin polymerization defects in OCI-AML3 cells upon treatment with cytochalasin D and latrunculin B. A single z slice is presented. C. Same as A, for the nanotube inhibitor L-778,123. D. Same as A, for the endoplasmic reticulum (ER)-Golgi transport inhibitor Brefeldin A. E. Same as A, for the mitochondrial reactive oxygen species (ROS) generator Rotenone.

**Figure S4.**
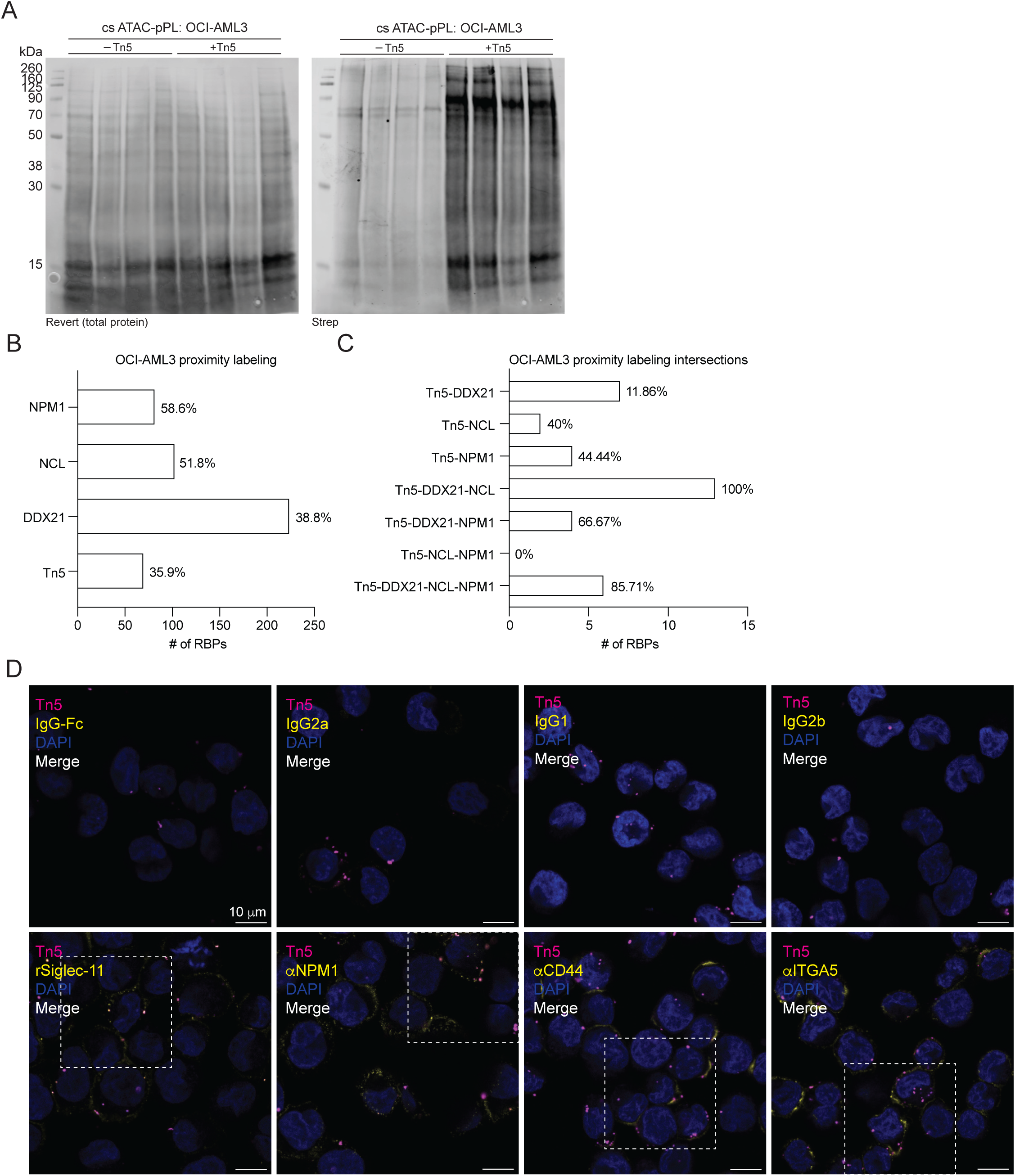
Characterization and quality control of csATAC-pPL. A. Western blot analysis of lysates from csATAC pPL reactions on OCI-AML3 cells with and without Tn5. Total protein (Revert stain) and biotinylated proteins (Strep) are presented for 4 biological replicates. B. Quantification of the fraction of RBP hits in OCI-AML3 proximity labeling datasets used for intersection analysis (csATAC pPL, ^44,45^). C. Quantification of the fraction of RBP hits in Tn5-containing overlaps between proximity labeling datasets. D. Uncropped confocal microscopy for co-localization analysis, including the corresponding isotype stains for each indicated antibody. The z slice used for co-localization analysis is shown.

**Figure S5.**
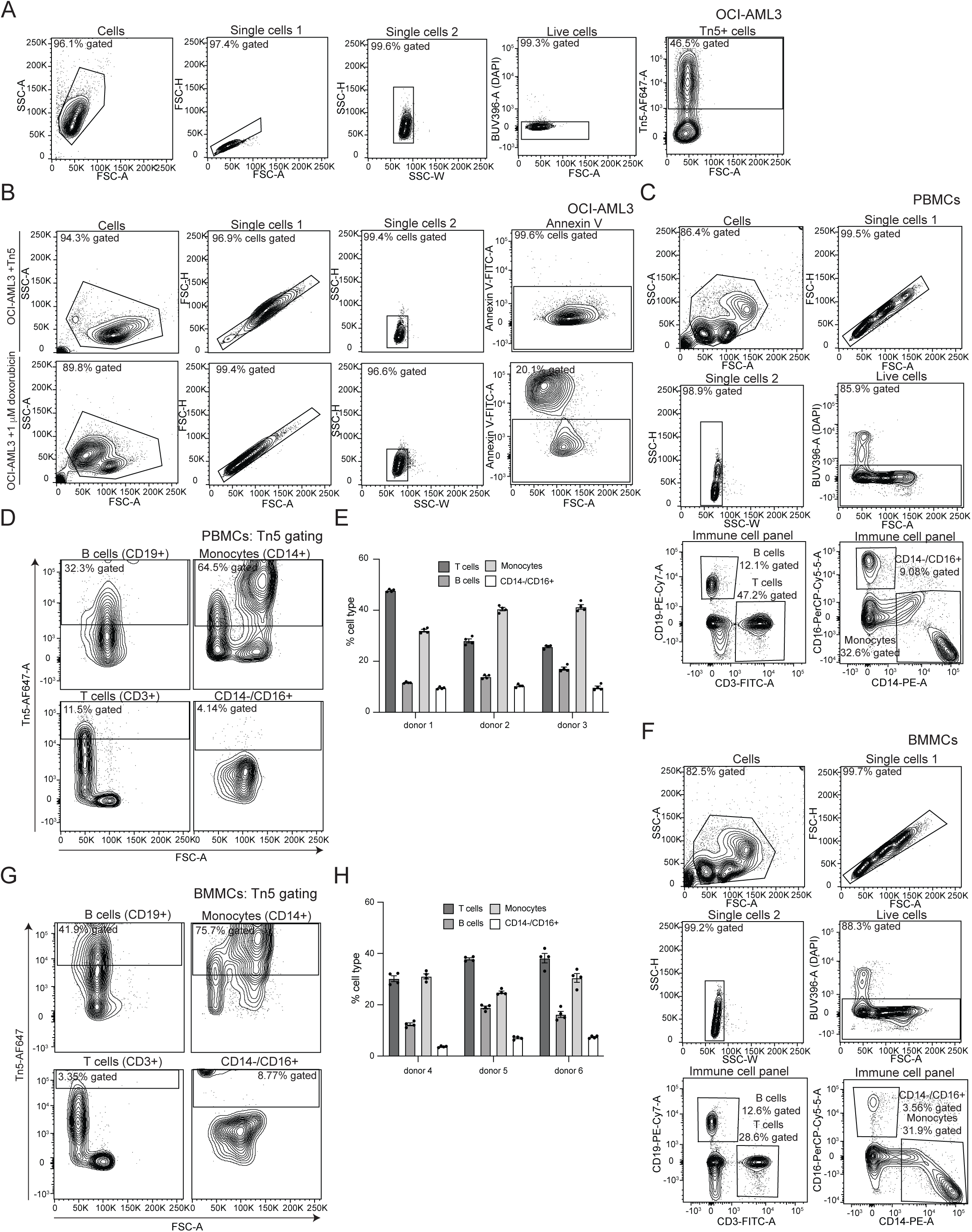
csATAC-See flow cytometry on OCI-AML3 cells, human PBMCs, and human BMMCs. A. Gating scheme for csATAC-See flow cytometry on OCI-AML3 cells. B. Gating scheme for Annexin V staining on OCI-AML3 cells. C. Gating scheme for PBMCs stained with an immune cell antibody panel. D. Tn5 gating on PBMCs for each cell type. E. Quantification of the fraction of each immune cell type for each donor (n= 4 technical replicates per donor: ±Tn5, ±rDNASE1L3). F. Gating scheme for BMMCs stained with an immune cell antibody panel. G. Tn5 gating on BMMCs for each cell type. H. Quantification of the fraction of each immune cell type for each donor (n= 4 technical replicates per donor: ±Tn5, ±DNASE1L3).

**Figure S6.**
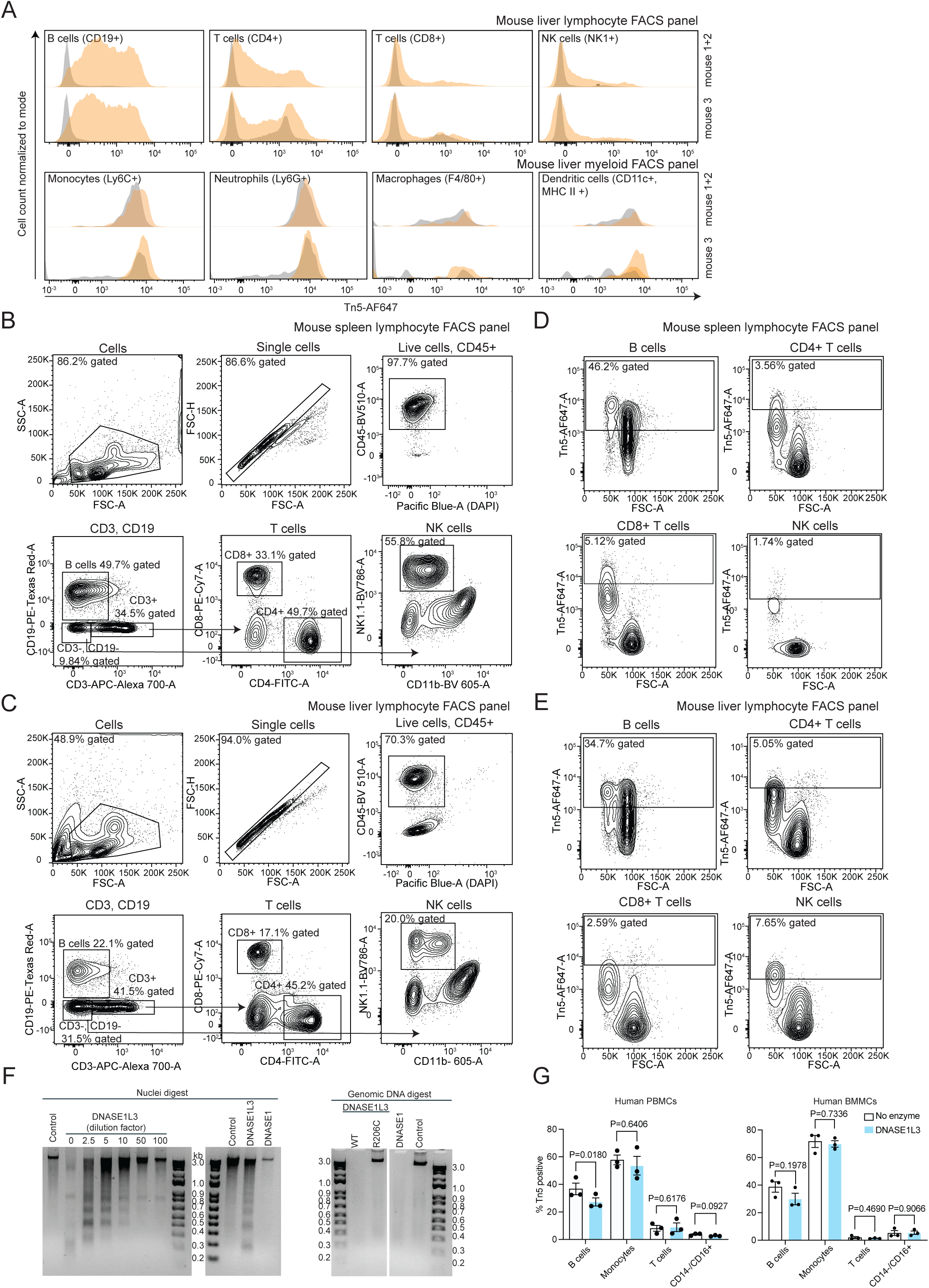
Cell surface DNA on B cells is sensitive to recombinant DNASE1L3. A. Live cell flow cytometry of mouse liver cells using a lymphocyte and myeloid antibody panel. B. Gating scheme for mouse spleen cells stained with a lymphocyte antibody panel. C. Gating scheme for mouse liver cells stained with a lymphocyte antibody panel. D. Tn5 gating on mouse spleen cells for each lymphocyte cell type. E. Tn5 gating on mouse liver cells for each lymphocyte cell type. F. *In vitro* nuclei and genomic DNA digestion with recombinant DNASE1L3 (wild type and R206C/vehicle) or DNASE1. G. Quantification of PBMCs and BMMCs gated as Tn5 positive for each cell type, with or without pre-treatment with recombinant DNASE1L3. *P* values were calculated using an unpaired student’s t-test (n= 3 donors).

**Figure S7.**
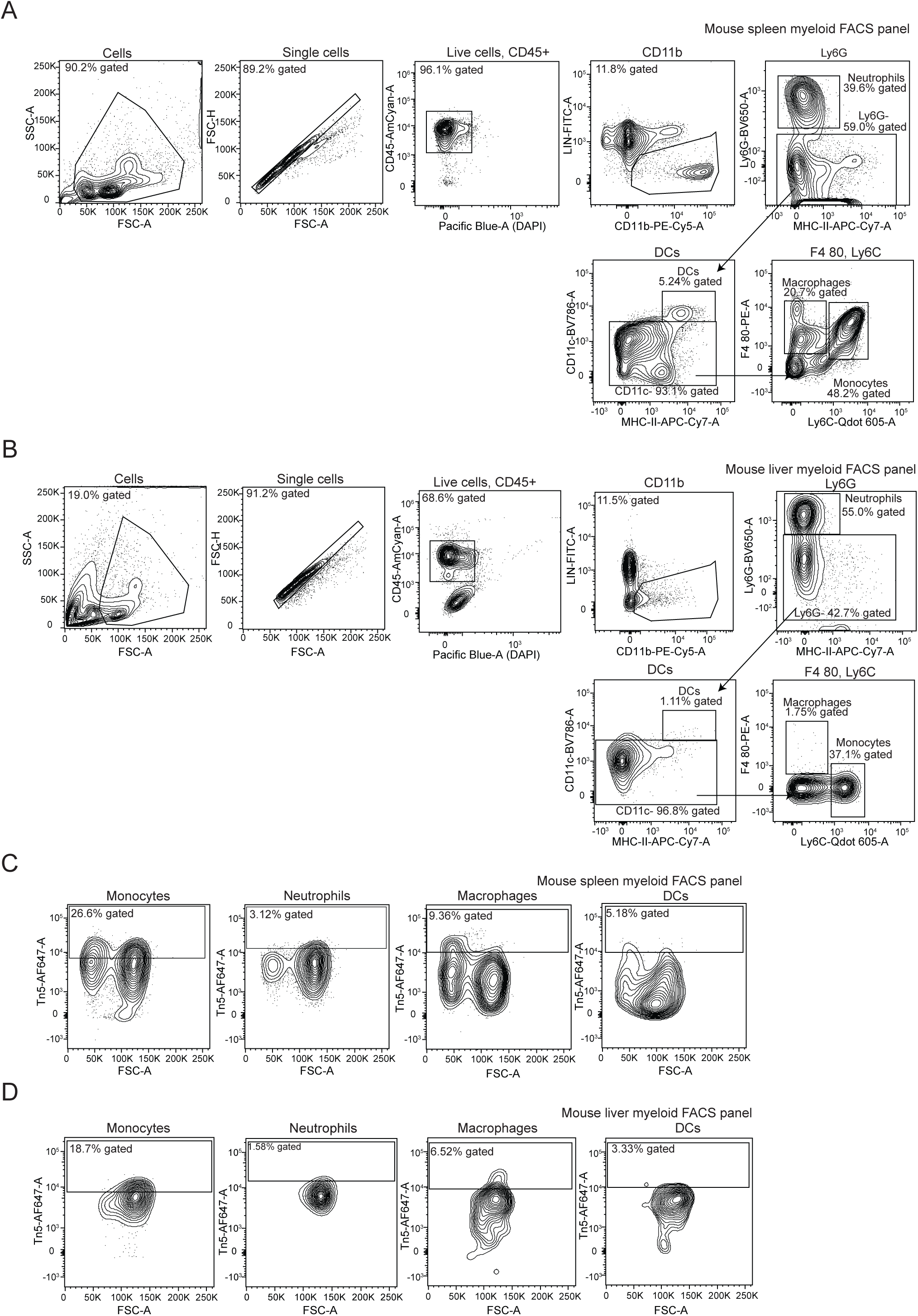
csATAC-See flow cytometry on mouse spleen and liver cells stained with a myeloid antibody panel. A. Gating scheme for mouse spleen cells stained with a myeloid antibody panel. B. Gating scheme for mouse liver cells stained with a myeloid antibody panel. C. Tn5 gating on mouse spleen cells for each myeloid cell type. D. Tn5 gating on mouse liver cells for each myeloid cell type.

## Table Legends

**Supplementary Table 1. MRM transitions used for quantification of nucleosides.**

**Supplementary Table 2. csDNA-proximal proteins identified by csATAC-pPL MS.**

**Supplementary Table 3. Peaks identified in standard and csATAC-Seq.**

**Supplementary Table 4. Oligo sequences.**

**Supplementary movie 1. 3D reconstruction of OCI-AML3 cells labeled with Tn5.**

## Data availability

Nucleoside and protein mass spectrometry data have been deposited on MassIVE (MSV000098850, MSV000098858) and sequencing data have been deposited in the NCBI SRA (PRJNA1310645).

## Methods

### Mammalian cell culture

All cells were cultured in 5% CO_2_ at 37°C and maintained as mycoplasma negative. Adherent cell lines were split when cells reached 80-90% confluency by rinsing cells with 1x PBS and treating cells with Trypsin (Quality Biological) or TrypLE (Thermo) for 5 minutes at 37°C, and quenched with the addition of complete media. A549 (ATCC) and HEK293T (ATCC) adherent cells were grown in 1x DMEM (ThermoFisher Scientific) supplemented with 10% heat inactivated fetal bovine serum (FBS, ThermoFisher Scientific) and 1% Penicillin and Streptomycin (Pen/Strep, ThermoFisher Scientific). OCI-AML3 (ATCC) suspension cells were grown in 1x RPMI-1640 supplemented with 10% heat inactivated FBS and 1% Pen/Strep and split when cells reached a density of 2 million cells per mL by resuspending cell pellets in fresh media. Primary human peripheral blood mononuclear cells (PBMCs) and bone marrow mononuclear cells (BMMCs) (Stem Cell) were thawed into 1x RPMI-1640 supplemented with 10% FBS and 1% Pen/Strep and spun at 300xg for 10 minutes at room temperature, followed by another wash with complete media.

Whole blood was collected in EDTA tubes and PBMCs were isolated by gently layering blood over a 15 mL cushion of Ficoll-Paque Plus (Global Life Sciences Solutions) and spinning cells at 500xg for 30 minutes at 4°C, with the brake turned off. The top plasma layer was discarded and the intermediate layer containing Ficoll and PBMCs was removed to a fresh tube with 20 mL 1x PBS, followed by centrifugation at 500xg for 10 minutes at 4°C to pellet PBMCs. B cells were isolated with a pan B cell isolation kit (Miltenyi Biotech).

### Mouse strains

6–8 week-old female and male wild-type (C57BL/6J, Jax #000664) mice were purchased from The Jackson Laboratory. Mice were housed in pathogen-free conditions at the department of Animal Resources at Children’s Hospital (ARCH) in Boston Children’s Hospital. All procedures were approved under the Institutional Animal Care and Use Committee (IACUC).

### Human Subjects and Sample Acquisition

Cryopreserved human peripheral blood mononuclear cells (PBMCs) were acquired from The Genotype and Phenotype (GaP) Registry at The Feinstein Institutes for Medical Research, Manhasset, New York, USA. Samples were selected based on homozygosity for the R206C (rs35677470) polymorphism: either homozygous for the reference allele (206RR, rs35677470 gg) or homozygous for the minor/disease risk allele (206CC, rs35677470 aa). These PBMCs were sourced from a combination of a retrospective sample bank (as detailed in ^60^) and GaP enrollment collections.

### Ethics Approval for Human Subjects

All GaP Registry samples were collected from consented participants under an Institutional Review Board (IRB)-approved protocol (IRB# 09–081) at The Feinstein Institutes for Medical Research. Samples were de-identified (pseudonymized) before release for research. This study was approved by The Committee for Participant Protection at The Feinstein Institutes for Medical Research (Protocol ID: TAP0307.5.85).

### Trypsin release, DNA extraction, enzyme digestions, and Southern blotting

Cells were treated with Trypsin (Quality Biological) or TrypLE (ThermoFisher Scientific) (4 mL Trypsin/TrypLE per 15 cm plate of adherent cells, 1 mL Trypsin/TrypLE per 1 million suspension cells) for 10 minutes at 37°C and the resulting Trypsin-cell mixture was spun down twice at 400xg for 3 minutes at room temperature to pellet and remove cells. Trypsin was lyophilized and resuspended in 1 mL nuclease-free water, followed by the addition of 2 mL (2 volumes) DNA binding buffer (Zymo Research) and 6 mL (2 volumes water + DNA binding buffer) 100% ethanol, vortexing between each addition. Samples were purified over Zymo-Spin III columns (Zymo Research) and washed 3 times: once with 400 µL RNA Prep buffer (3M guanidinium hydrochloride in 80% ethanol), and twice with 400 µL 80% ethanol. 60 µL 2 mg/mL Proteinase K (NEB) was directly applied to the column membrane and incubated for 45 minutes at 37°C, followed by the addition of 60 µL water and a 30 second spin at 10,000xg to elute DNA. Samples were further purified to get rid of Proteinase K by an additional round of purification over Zymo-Spin III columns. For whole cell DNA isolation, cells were lysed in DNA lysis buffer (Zymo Research) and lysates were directly applied to Zymo-Spin I columns, washed twice with 400 µL 80% ethanol, and eluted with water. All *in vitro* enzyme digestions were carried out in 10 µL reactions for 1 hour at 37°C in 1X TURBO DNase buffer (ThermoFisher Scientific) with 1 µL TURBO DNase (ThermoFisher Scientific), 1 µL Proteinase K, or 1 µL each of RNase A/T1 cocktail (ThermoFisher Scientific) and ShortCut RNase III (NEB). For Southern blots, DNA was digested with ClaI or NdeI in a 20 µL reaction containing 1x CutSmart buffer (NEB) for 2-3 hours at 37°C. Samples were resolved on a 0.7% agarose gel run at 2-3 V/cm for 3-6 hours. The gel was washed with 0.25M HCl in water for 10 minutes, denatured in 0.5M NaOH, 1.5M NaCl for 30 minutes, and neutralized with 1M Tris pH 7.0, 1.5M NaCl for 30 minutes, followed by downward transfer onto a Hybond N+ membrane (Cytiva) in 10X SSC (0.15M sodium citrate, 1.5M NaCl, pH 7.0) overnight at room temperature. The membrane was UV crosslinked and hybridized in 15 mL ULTRAhyb Buffer (ThermoFisher Scientific) with 90 µL denatured salmon sperm DNA (ThermoFisher Scientific) at 42°C with an ɑ- ^32^P-labeled 7kb mtDNA for 3-4 hours. The membrane was washed twice with 2X SSC, 0.1% SDS for 15 minutes each at room temperature, twice with 0.1X SSC, 0.1% SDS for 15 minutes each at 42°C, and exposed with a PhoshorScreen for imaging.

### DNA sequencing and analysis

DNA libraries were prepared from 50 ng RNase A/T1/III-treated whole cell and trypsin-released DNA in biological triplicate using the NEBNext Ultra II DNA Library Prep kit (NEB), with DNA fragmented into 200-450 base pair fragments. Libraries were sequenced using 100bp paired end reads (and 50bp paired end reads for whole cell and trypsin-released OCI-AML3 cells) on a NextSeq 550 at the Tufts genomics core facility. Reads were aligned to the GRCh38 reference genome using Bowtie2 ^75^. To calculate normalized mapped reads, the total number of mapped reads per chromosome were divided by the total number of reads per sample and chromosome length.

### LC–MS/MS Analysis of 8-oxo-dG in DNA Samples

DNA samples were enzymatically hydrolyzed into nucleosides using the following previous established protocols ^52^. Briefly, 1.0 U of Super Nuclease and 0.02 U of phosphodiesterase I (PDE-I) were added to the DNA solution in a digestion buffer containing 1 mM ZnCl₂ and 30 mM sodium acetate (pH 7.5), followed by overnight incubation at 37 °C. Subsequently, 2.0 U of recombinant shrimp alkaline phosphatase (rSAP) was added and incubated at 37 °C for 1 hour to ensure complete dephosphorylation of nucleotides to nucleosides. The digested samples were diluted with 400 μL of 0.1% trifluoroacetic acid (TFA) in water and purified using porous graphitized carbon (PGC) solid-phase extraction (Thermo Scientific). The cartridges were washed with 0.1% TFA in water and followed by elution with 40% acetonitrile containing 0.05% TFA and 80% acetonitrile containing 0.1% TFA. The eluates were combined, dried under vacuum using a CentriVap centrifugal concentrator (Labconco), and reconstituted in 0.1% formic acid (FA) in water for subsequent LC–MS/MS analysis.

Chromatographic separation was performed using a CORTECS T3 column (2.1 mm × 100 mm, 2.7 μm particle size, 120 Å pore size; Waters) on a Shimadzu Nexera X3 UHPLC system, consisting of LC-40D X3 binary pumps, a SIL-40C X3 autosampler, a CTO-40C column oven (maintained at 40 °C), and an SCL-40 system controller. The mobile phase consisted of 0.1% formic acid in water (buffer A) and 0.1% formic acid in acetonitrile (buffer B), with a flow rate of 0.3 mL/min. The gradient elution program was: 0–3 min, 0% B; 3–8 min, 0–6% B; 8–11.5 min, 6–60% B; 11.5–12.5 min, hold at 60% B; 12.5–12.6 min, return to 0% B; 12.6–15 min, re-equilibration at 0% B.

Mass spectrometric detection was performed on a SCIEX QTRAP 6500+ triple quadrupole mass spectrometer equipped with an ESI Turbo V source, operated in positive ion mode under a scheduled multiple reaction monitoring (MRM) method. The source parameters were set as follows: IonSpray voltage at 5000 V, source temperature at 500 °C, curtain gas at 35 psi, ion source gas 1 (GS1) and gas 2 (GS2) both at 55 psi. Compound-specific parameters included a declustering potential (DP) of 80 V, entrance potential (EP) of 10 V, collision energy (CE) of 30 V, and collision cell exit potential (CXP) of 10 V. Each transition was monitored with a dwell time of 100 ms. The monitored transitions as summarized in Table S1.

For quantification, data were acquired and processed in NumoFinder ^52^ and Skyline ^76^. The peak corresponding to 8-oxo-dG in DNA samples was identified based on co-elution with the synthetic 8-oxo-dG standard (Macklin) and confirmed by matching fragmentation patterns. Quantitative analysis was performed using extracted ion chromatograms and peak areas. Relative abundance of 8-oxo-dG was calculated by normalizing to deoxyguanosine.

### Sample preparation for cell surface ATAC-See and confocal microscopy

Equimolar amounts of 100 µM Tn5MErev and Tn5ME-A-AF647, and in a separate tube Tn5MErev and Tn5ME-B-AF647, were mixed and heated at 95°C for 5 minutes, then slow cooled to room temperature in a thermocycler to create double stranded oligonucleotides. A 25 µL Tn5 transposition reaction was assembled by mixing 3.125 µL of each annealed oligonucleotide (final concentration of 50 µM for each double stranded oligonucleotide), 12.5 µL 80% glycerol, 3 µL 2X dialysis buffer (100 mM HEPES pH 7.4, 0.2M NaCl, 0.2 mM EDTA, 2 mM DTT, 20% glycerol), and 3.33 µL Tn5 (Diagenode) (final Tn5 concentration of 5 µM) and incubating the reaction for 1 hour at room temperature, away from light (the transposition reaction can be made in advance and stored at -20°C). Cells (750,000 suspension cells or 150,000 adherent cells seeded onto a cover slip in a 12-well plate the night before) were washed with 1x PBS, then resuspended in a 50 µL reaction containing 2.5 µL pre-assembled transposition reaction and 25 µL 2X TD buffer (20 mM Tris pH 7.6, 4 mM MgCl_2_) and incubated for 5 minutes at 37°C. The reaction was quenched twice with 1 mL ice cold FACS buffer (0.5% BSA in 1x PBS), followed by a 15 minute fixation at room temperature with 4% formaldehyde and mounting on glass slides with 50% glycerol in 1x PBS containing 1x DAPI. Suspension cells were applied to glass slides by spinning cells for 3 minutes at 500xg on a CytoSpin 1867. For samples that were co-stained with antibodies, after Tn5 labeling and washing with FACS buffer, cells were blocked with 3.75 µL human TruStain FcX (Biolegend) in 750 µL FACS buffer for 15 minutes on ice. Cells were then stained with 2.5 µg/mL unconjugated primary antibody (mouse isotype (Biolegend), NPM1 ^45^, CD44 (Biolegend), CD49e/ITGA5 (Biolegend), ꞵ2 microglobulin (Santa Cruz)) for 45 minutes on ice, washed twice with FACS buffer, stained with 2.5 µg/mL goat anti-mouse AF488 Plus (Invitrogen) for 30 minutes on ice, washed twice with FACS buffer, then fixed and applied to glass slides as noted above. For Siglec-11, 1 µg/mL rSiglec-11-Fc (R&D Systems) or IgG-Fc was pre-complexed with 0.5 µg/mL goat anti-human AF488 (Invitrogen) in FACS buffer for 45 minutes on ice, then incubated with Tn5 labeled cells for 30 minutes on ice, followed by fixation and mounting.

### Confocal microscopy data acquisition and analysis

Cells were imaged on a Leica TCS SP8 STED ONE microscope with a 63x oil immersion objective across 6 x µm z-slices (or 12 x 0.5 µm z-slices for 3D volumetric reconstruction) using Leica’s line-sequential scanning method and 1024 by 1024 resolution with a pinhole size of 1 AU. DAPI signal was collected with a PMT detector and all other channels were collected on a Hybrid detector. Images were taken from biological triplicates. For co-staining experiments, cells were imaged on a Zeiss LSM 900 microscope with a 63x oil immersion and acquired with an Airyscan 2 detector. 3 regions of interest (ROIs) were collected for each antibody. Spot number and intensities were measured in 3D from each image using Fiji. For co-localization, image processing was done in Imaris Microscopy Image Analysis software (Oxford Instruments) and co-localization analysis was performed as in ^44^. Briefly, a single z stack in the middle of the cells was selected and the spot-finder function was used to identify spots by thresholding spot quality at the elbow of the distribution, resulting in a series of x- and y-positions for each spot from each channel. A custom Python script (https://github.com/FlynnLab/jonperr) was used to identify nearest neighbors of each spot with a k-d tree algorithm (scipy.spatial.KDTree). A Manders’ colocalization coefficient was calculated to determine the relative fraction of each spot type (channel 1) within the other pair’s spots (channel 2), in both directions.

### Sample preparation for electron microscopy

Cells (750,000 suspension cells or 150,000 adherent cells seeded onto a cover slip in a 12-well plate the night before) were washed with 1x PBS, then resuspended in a 50 µL reaction containing 2.5 µL pre-assembled transposition reaction made with Tn5ME-A-biotin and Tn5ME-B-biotin and 25 µL 2X TD buffer (20 mM Tris pH 7.6, 4 mM MgCl_2_) and incubated for 5 minutes at 37°C (transmission electron microscopy, TEM) or 20 minutes at 37°C (scanning electron microscopy, SEM), then washed twice with 1 mL ice cold FACS buffer. Cells were resuspended in Streptavidin Gold, 15 nm particles (Fisher Scientific) diluted 1:5 in FACS buffer, incubated for 45 minutes on ice, and washed twice with 1x PBS. Cells were fixed in 2.5% glutaraldehyde, 0.1 M cacodylate buffer pH 7.4 for 1 hour at room temperature. For TEM, cells were cryo-sectioned and imaged on a Tecnai G^2^ Spirit BioTWIN microscopy under 9300x direct magnification at the Harvard Medical School Electron Microscopy Facility. SEM sample preparation and imaging were performed by the Harvard Medical School Electron Microscopy Facility as per ^77^ using 70k zoom.

### Super resolution sample preparation, imaging, and reconstruction

150,000 A549 cells were seeded onto a cover slip in a 12-well plate the night before, then washed with 1x PBS and labeled for csATAC-See for 5 minutes at 37°C, washed twice with 1 mL ice cold FACS buffer, and sequentially stained with 2.5 µg/mL ꞵ2 microglobulin (Santa Cruz) and 2.5 µg/mL goat anti-mouse AF568 (Invitrogen), as above. Cells were washed once with 1x PBS and fixed with 4% formaldehyde, 0.2% glutaraldehyde for 15 minutes at room temperature, away from light, and stored in 1x PBS until imaging.

Image stacks of single molecules were obtained on a custom optical setup consisting of an inverted microscope (Nikon Instruments, Eclipse Ti2) with Perfect Focus System. Fluorescence was collected via a high NA objective (Nikon Instruments, Apo SR TIRF 100, NA 1.49, Oil). 1 W 560- and 642 nm lasers (MPB Communications, 1 W) were used for excitation. The power of the laser beam was controlled in free space using a filter wheel (Thorlabs, FW212CNEB). The laser beam was passed through a cleanup filter (Chroma Technology, ZET561/10) and coupled into the microscope objective using a beam splitter (Chroma Technology, ZT561RDC). Fluorescence was spectrally filtered with an emission filter (Chroma Technology, ET600/50m, and ET575LP) and imaged on an sCMOS camera (Hamamatsu Orca Fusion) without further magnification, resulting in an effective pixel size of 130 nm after 2×2 binning. The center of the camera was used as the region of interest. Raw microscopy data was acquired using μManager (Version 2.0.3).

A reducing oxygen scavenging buffer that induces blinking of single fluorophores was employed according to ^78^. The blinking buffer consisted of 2 μL/mL catalase (Sigma-Aldrich), 10% (w/v) glucose (BD Biosciences), 100 mM Tris-HCl (Thermo Fisher), 560 μg/mL glucose oxidase (Sigma-Aldrich), and 20 mM cysteamine (Sigma-Aldrich). First, DL imaging was performed with low-intensity illumination of a few W/cm^2^ at the respective excitation wavelength. Then, the laser power was increased to approx. 3 kW/cm^2^. Image acquisition was started after a short delay in order to ensure that most fluorophores were shelved into a dark state. 20000 frames were acquired with a frame time of 50 ms. The two channels were registered by imaging TetraSpeck beads (Fisher Scientific) in both channels, followed by affine transformation.

### ATAC-Seq sample preparation and analysis

csATAC-Seq was performed as per confocal imaging on 1 million OCI-AML3 cells per sample for 20 minutes at 37°C using AF647-labeled oligonucleotides. Following washes with FACS buffer, cells were resuspended in 1 mL Trypsin and incubated for 10 minutes at 37°C. Cells were pelleted twice at 400xg for 3 minutes and Trypsin was collected, lyophilized, and DNA was extracted as above. Whole cell ATAC-Seq was performed as per ^79,80^. Briefly, a 25 µL Tn5 transposition reaction was assembled by mixing 3.125 µL of each annealed oligonucleotide (final concentration of 50 µM for each double stranded oligonucleotide), 12.5 µL 80% glycerol, 3 µL 2X dialysis buffer (100 mM HEPES pH 7.4, 0.2M NaCl, 0.2 mM EDTA, 2 mM DTT, 0.2% Triton X-100, 20% glycerol), and 3.33 µL Tn5 (Diagenode) (final Tn5 concentration of 5 µM) and incubating the reaction for 1 hour at room temperature, away from light. 50,000 OCI-AML3 cells per sample were resuspended in 50 μL ice cold lysis buffer (10 mM Tris pH 7.5, 10 mM NaCl, 3 mM MgCl_2_, 0.1% Igepal, 0.1% Tween-20, 0.01% Digitonin) and incubated on ice for 3 minutes before the addition of 1 mL ice cold wash buffer (10 mM Tris pH 7.5, 10 mM NaCl, 3 mM MgCl_2_, 0.1% Tween-20) and mixing by inversion. Samples were centrifuged for 5 minutes at 500xg and 4°C to pellet nuclei, and the nuclear pellet was resuspended in 50 μL reaction containing 2.5 µL pre-assembled transposition reaction and 25 µL 2X TD buffer (20 mM Tris pH 7.6, 4 mM MgCl_2_, 20% dimethylformamide) and incubated for 20 minutes at 37°C. The reaction was quenched with the addition of 1 mL genomic DNA lysis buffer (Zymo Research) and purified using a Zymo-Spin I column.

For cell surface and whole cell ATAC-Seq, eluted DNA was resuspended in 20 µL water and mixed with 0.5 µL100 µM forward and reverse Nextera PCR primers, 0.35 µL 25x Sybr Green, and 20 µL NEBNext Ultra II Q5 Master Mix (NEB). Reactions were incubated at 72°C for 5 minutes, then 12-15 cycles of 10 seconds at 98°C, 30 seconds at 63°C, and 1 minute at 72°C. Libraries were purified using AMPure XP beads (Beckman Coulter) and sequenced with 100bp paired end reads on a NextSeq 550 at the Tufts genomics core facility. Reads were aligned to the GRCh38 reference genome using BWA ^81^ and analyzed using a custom ATAC-Seq pipeline from Basepair that retained mitochondrial-mapped reads. Peaks were called using MACS2 ^82^.

### Inhibitors and live cell enzyme treatments

For live cell benzonase treatments, 750,000 OCI-AML3 cells were resuspended in 500 µL pre-warmed FACS buffer with 2 mM MgCl_2_ and 2 µL benzonase nuclease (Millipore Sigma) and incubated for 30 minutes at 37°C. Live cell RNase treatments were performed in 500 µL pre-warmed FACS buffer with 2 mM MgCl_2_, 1 µL RNase A/T1 cocktail, and 1 µL ShortCut RNase III for 30 minutes at 37°C. A pool of heparinases I, II, and III (NEB) was added directly to cell media at a concentration of 1:1000 and incubated for 30 minutes at 37°C. 100 ng/mL ethidium bromide was added directly to cell cultures for 96 hours, passaging cells once in between and resuspending in fresh media containing 100 ng/mL ethidium bromide. Inhibitors were added directly to cell media at the following concentrations and times, all at 37°C: 25 µM CCCP (TargetMol) for 1 hour, 1 µM Cytochalasin D (Cayman Chemical Company) for 2 hours, 10 nM Latrunculin B (Sigma-Aldrich) for 2 hours, 10 µM L-778,123 (Cayman Chemical Company) for 2 hours, 5 µg/mL Brefeldin A (Sigma-Aldrich) for 2 hours, and 1 µM Rotenone (Sigma-Aldrich) for 1 hour. Phalloidin staining to validate actin inhibitors was performed by fixing 750,000 cells with 4% formaldehyde for 15 minutes at room temperature, followed by permeabilization with 0.1% Triton X-100 for 3 minutes at room temperature, washing twice with 1x PBS, and resuspending cells in 150 µL 1x Phalloidin-iFluor 488 conjugate (abcam) for 30 minutes at room temperature away from light. Cells were washed twice with 1x PBS, cytospun onto glass slides, and mounted with DAPI. Recombinant human full-length DNASE1L3 enzymes (wild-type and R206C variant), each engineered with an internal FLAG tag as described by ^65^, were custom-generated by GenScript (Piscataway, NJ, USA). These proteins were expressed in TurboCHO cells, and attempts were made to purify both variants from cell culture supernatants via anti-FLAG affinity chromatography. However, the R206C variant, which is known to be secretion defective, was not secreted into the supernatant. Consequently, its purification eluate contained no detectable protein or enzyme activity and was utilized as a vehicle control in this study. Live cell treatment with recombinant DNASE1L3 was performed on 1 million cells in 180 µL FACS buffer containing 1 mM MgCl_2_, 1 mM CaCl_2_, and 20 µL 5 ng/µL DNASE1L3 for 30 minutes at 37°C. Live cell treatment of isolated B cells was performed on 250,000 cells in 50 µL FACS buffer with 1 mM MgCl_2_, 1 mM CaCl_2_, and 5 µL 5 ng/µL DNASE1L3 or 0.5 µL 500 ng/µL recombinant human DNASE1 (Cusabio) for 30 minutes at 37°C. *In vitro* digests on purified nuclei and genomic DNA were performed as per ^60^.

### Cell surface protein proximity labeling, peptide generation, mass spectrometry, and data analysis

5 million OCI-AML3 cells per biological replicate were washed once with 1x PBS, resuspended in a 50 µL reaction containing 2.5 µL pre-assembled transposition reaction made with Tn5ME-A-biotin and Tn5ME-B-biotin and 25 µL 2X TD buffer (20 mM Tris pH 7.6, 4 mM MgCl_2_) and incubated for 20 minutes at 37°C. Following 2 washes with 1 mL ice cold FACS buffer, cells were resuspended in 500 µL FACS buffer with 2.5 µg Streptavidin-HRP (Thermo Scientific) and incubated for 45 minutes on ice in the dark, inverting to mix every 15 minutes. Cells were centrifuged at 400xg for 3 minutes at 4°C, washed once with ice cold 1x PBS, resuspended in 990 µL 100 µM biotin-tyramide (ApexBio) with 10 µL 100 mM H_2_O_2_, and incubated for 100 seconds with occasional inversion to mix. The labeling reaction was quenched with 250 µL FACS buffer containing 50 mM sodium ascorbate and 25 mM sodium azide, washed once with ice cold 1x PBS, and lysed in CLIP lysis buffer (50 mM Tris (pH 7.5), 1% IGEPAL, 0.5% Na-deoxycholate, 150 mM NaCl, 5 mM EDTA). Lysates were reduced with 1% SDS and 5 mM DTT in CLIP lysis buffer for 30 minutes at 65°C, then alkylated with 25 mM iodoacetamide (Sigma-Aldrich) for 30 minutes at room temperature. Biotinylated proteins were immunoprecipitated with neutravidin beads (Thermo Scientific) and tryptic peptides were generated and processed as per ^44^. Briefly, beads were washed twice with 1 mL 4 M NaCl with 100 mM HEPES, twice with 2 M guanidinium hydrochloride with 100 mM HEPES, twice with LC-MS grade water, and once with 50 mM ammonium bicarbonate. Samples were digested at 37°C overnight in 200 µL 50 mM ammonium bicarbonate with 1 µg trypsin (Promega) and beads were rinsed with 400 µL 50 mM ammonium bicarbonate and 400 µL 50% acetonitrile (in water), combining both washes with the trypsin eluate. The pooled eluate was spiked with 0.1% formic acid (Fisher Scientific), lyophilized, and peptides were desalted with C18 spin columns (Thermo Scientific).

An LC-MS/MS system consisting of a Vanquish Neo UHPLC coupled to an Orbitrap Exploris 240 or Ascend (Thermo Scientific) was used for peptide analysis. Separation of peptides was carried out on an Easy-Spray^TM^ PepMap^TM^ Neo nano-column (2 µm, C18, 75 µm x 150 mm) at room temperature with a mobile phase. The chromatography conditions consisted of a linear gradient from 2 to 32% solvent B (0.1% formic acid in 100% acetonitrile) in solvent A (0.1% formic acid in water) over 75 minutes and then 45 to 98% solvent B over 15 minutes at a flow rate of 300 nL/min.

The mass spectrometer was programmed for data-dependent acquisition (DDA) mode. The MS1 was acquired over a mass range of 295-1100 m/z with a resolution of 120,000 an AGC target of 300%, and a maximum injection time of 50 ms. For the MS2, the top 20 most intense ions were selected for MS/MS by high-energy collision dissociation (HCD) at 30 NCE, with a resolution of 15,000, AGC target of 75%, and maximum injection time of 60 ms. Raw files were processed with FragPipe ^83^. Carbamidomethyl (C) was set as a static modification while oxidation (M) and acetylation (protein N-terminus) were selected as variable modifications.

Data was filtered to remove contaminants and include proteins with values in at least 3 out of 4 biological replicates for samples labeled with Tn5. Filtered data was uploaded to Perseus (MaxQuant), log transformed and underwent imputation using normalization to adjust for values of 0. Significance was determined by performing a Welch’s t-test (S0=1.5) on non log-transformed data.

### Flow cytometry and cell staining

Annexin V staining was performed using the Annexin V FITC kit from BD Biosciences, following the manufacturer’s protocol. For csATAC-See on immune cells, 1 million frozen PBMCs or BMMCs were blocked in 100 µL FACS buffer containing 12.5 pmol single-stranded, non-targeting DNA oligos for 5 minutes at 37°C, then diluted with 400 µL FACS buffer and centrifuged at 400xg for 5 minutes at room temperature. Cells were resuspended in 50 µL transposition reactions with or without Tn5, as described above, and incubated for 5 minutes at 37°C, followed by 2 washes with ice cold FACS buffer. Cells were Fc blocked for 15 minutes on ice using 5 µL Human TruStain FcX (Biolegend) in 1 mL FACS buffer per million cells, stained with their respective antibody panels (500,000 cells per panel, using an antibody dilution of 1:100) for 30 minutes on ice before resuspension in 400 µL FACS buffer with 1x DAPI.

To process mouse organs, spleens and livers were smashed over 70 µm filters with 45 mL 1x PBS and centrifuged at 1500 rpm for 5 minutes. Spleens were manually resuspended and red blood cell lysis was performed using 1x ACK (Thermo Scientific) for 5 minutes on ice, then washed with PBS. Livers were resuspended in 1 mL 42% Percoll, spun at 800xg for 20 minutes with no brake, manually resuspended and red blood cell lysis was performed as above. Single cell suspensions containing 10 million cells were blocked in 1 mL FACS buffer with 125 pmol single-stranded, non-targeting DNA oligos for 5 minutes at 37°C, resuspended in 100 µL transposition buffer with or without Tn5 for 5 minutes at 37°C, and washed twice with ice cold FACS buffer. Cells were stained with Zombie violet for 10 minutes at room temperature, washed with 1x PBS, Fc blocked for 15 minutes on ice using 10 µL anti-mouse CD16/32 TruStain FcX (Biolegend) in 100 µL FACS buffer, and stained with their respective antibody panels (5 million cells per panel, using an antibody dilution of 1:100) for 30 minutes on ice before resuspension in 400 µL FACS buffer. All data was acquired on a Fortessa cytometer and analyzed using FlowJo.

## Author Contributions

R.A.F. and J.P. conceived the project and designed experiments. J.P. performed most of the experiments and analysis. V.P. performed mouse work with supervision from I.Z. K.A. performed super resolution microscopy with supervision from L.M. B.M.G. designed and assisted with FACS experiments. F.N.L.V., L.Y., C.B.C., Y.Z., Y.X., and B.A.G. collected and analyzed mass spectrometry data. A.C. performed southern blot analysis with supervision from S.A. H.W., L.Z., and P.K.G. provided patient samples. K.R.S. designed DNASE1L3 experiments, provided GaP participant samples, and recombinant proteins. R.A.F, L.M., S.A., B.A.G., I.Z., and K.R.S. acquired funding. J.P. wrote the manuscript and J.P. and R.A.F. revised the manuscript with input from all authors.

## Funding

J.P. is supported by a postdoctoral fellowship from the Canadian Institutes for Health Research. This work was supported by grants from the Rita Allen Foundation (R.A.F.), the Scleroderma Research Foundation (R.A.F.), and the National Institutes of Health, National Institute of General Medical Sciences under award number R35GM151157 (R.A.F.), National Institute of Diabetes and Digestive and Kidney Diseases under award number R01DK107716, R.A.F. is the Bakewell Foundation-Rachleff Innovator of the Damon Runyon Cancer Research Foundation (DRR-74-23). K.A. and L.M. gratefully acknowledge financial support from the Else-Kröner-Fresenius-Stiftung (grant ID 2020_EKEA.91), the German Research Foundation (DFG, grant ID 529257351), and the Wilhelm-Sander-Stiftung (grant ID 2023.025.1), as well as by the Max Planck Society.

## Declaration of interests

R.A.F. is a stockholder of ORNA Therapeutics and a member of the board of directors and a stockholder of Blue Planet Systems. The other authors declare no competing interests.

## Acknowledgments

We thank Eliezer Calo, Sun Hur, John Pluvinage, Franck Barrat, Deepak Rao, Xu Zhou, Joyce Chang, Vijay Rathinam, Vincent Graziano, and Paul Hoover for helpful discussions. Electron microscopy sample preparation and imaging were performed in the HMS Electron Microscopy Facility. We thank the Boston Children’s Hospital IDDRC Cellular Imaging Core funded by NIH P50 HD10531 and the Boston Children’s Hospital IDDRC Molecular Genetics Core funded by NIH P30 HD 18655. DNA sequencing was performed at the Tufts University Core Facility. We thank Basepair for assistance with sequencing analysis.

